# SCALPEL: A pipeline for processing large-scale spatial transcriptomics data

**DOI:** 10.64898/2026.01.09.698732

**Authors:** Michael Kunst, Lindsey Ching, Jacob Quon, Rémi Mathieu, Madeleine N. Hewitt, Stephanie C Seeman, Angela Ayala, Emily C. Gelfand, Brian Long, Naomi X. Martin, Josh Nagra, Paul A. Olsen, Alana Oyama, Nasmil J. Valera Cuevas, Chelsea M. Pagan, Susan M. Sunkin, Jeanelle Ariza, Kimberly A. Smith, Delissa A. McMillen, Hongkui Zeng, Jack Waters

## Abstract

Spatial transcriptomics enables the precise mapping of gene expression patterns within tissue architecture, offering unprecedented insights into cellular interactions, tissue heterogeneity, and disease pathology that are unattainable with traditional transcriptomic approaches. We present a tool for processing spatial transcriptomics data, SCALPEL (Spatial Cell Analysis, Labeling, Processing, and Expression Linking). SCALPEL is specifically designed to support the analysis of large, atlas-level datasets. Our new workflow features advanced 3D segmentation optimized for dense and heterogeneous tissues, refined filtering criteria, and transcriptome-based doublet detection to remove low-quality or artifactual cells. Cell type label transfer from existing taxonomies is further improved through updated filtering thresholds. Spatial domain detection is incorporated to capture local transcriptomic organization, and tissue sections are registered to the Allen Mouse Brain Common Coordinate Framework version 3 (CCFv3) for precise anatomical alignment. Genome-wide expression imputation from single-cell RNA-sequencing (scRNAseq) further enriches the dataset. Crucially, we benchmark the performance of this updated pipeline against a previously published version of our whole-mouse-brain (WMB) dataset (Yao et al., 2023b), demonstrating substantial improvements in cell number, expression profile clarity, and spatial registration. These advances provide a robust foundation for downstream spatial analyses and set a new standard for large-scale spatial transcriptomics studies.

## Introduction

Single-cell spatial transcriptomics is a rapidly evolving field that provides unprecedented insights into the complex spatial organization of individual cells within their native tissue contexts (Close et al., 2021; Tian et al., 2023). This technology has the potential to revolutionize our understanding of cellular heterogeneity, cell-cell interactions, and the spatial patterns of gene expression that underpin tissue function and disease progression (Marx, 2021). With the availability of commercial products, the use of spatial transcriptomic methods is becoming more widely adopted by the scientific community as demonstrated by the ever-increasing number of datasets available on public repositories such as cellxgene (https://cellxgene.cziscience.com) or the Brain Image Library (BIL, https://www.brainimagelibrary.org/). In addition, there are ongoing projects aiming to map entire tissues such as BICAN (https://www.portal.brain-bican.org), HubMap (https://hubmapconsortium.org), and ImmGen (https://www.immgen.org). Standardized processing code is needed to compare datasets. Other tools to process spatial transcriptomics exist, but they focus on processing and analyzing individual sections rather than multi-section atlas datasets (Blampey et al., 2024; Cisar et al., 2023; Sun et al., 2025). In this paper, we present a suite of comprehensive processing code for single-cell spatial transcriptomics data acquired using subcellular spatial transcriptomics. The individual steps included are:

- Cell Segmentation
- Low quality cell filtering
- Doublet removal
- Mapping to reference taxonomy and additional filtering of low-quality mapping
- Spatial Domain detection
- Registration to CCF
- Gene imputation

Through the course of this paper, we will detail each step and discuss the rationale behind the choice of methods. We demonstrate the utility of our pipeline on a published and freely available Allen adult mouse brain dataset (Yao et al., 2023b).

## Results

### Workflow

We recently generated a whole mouse brain spatial transcriptomics dataset (Yao et al., 2023b), comprising 66 coronal sections collected at approximately 200 µm intervals (Fig. 1a). For this study, we imaged 500 genes selected to distinguish cell types identified in our scRNAseq whole-brain taxonomy, which was developed from profiling about 4 million cells (Fig. 1b, Table S1). After evaluating tissue quality with the MERQUACO tool (Martin et al., 2025), 59 high-quality sections remained for analysis Fig. 1c. Our data processing pipeline is structured in two main parts. The first part is an automated workflow that includes four key steps (Fig. 1d): (1) cell segmentation to assign mRNA molecules to individual cells, (2) removal of low-quality cells, (3) doublet detection and exclusion, and (4) mapping cells to our reference taxonomy followed by filtering out cells with low mapping confidence. This automated pipeline ensures standardized and reproducible preprocessing of the dataset.

**Figure 1.**
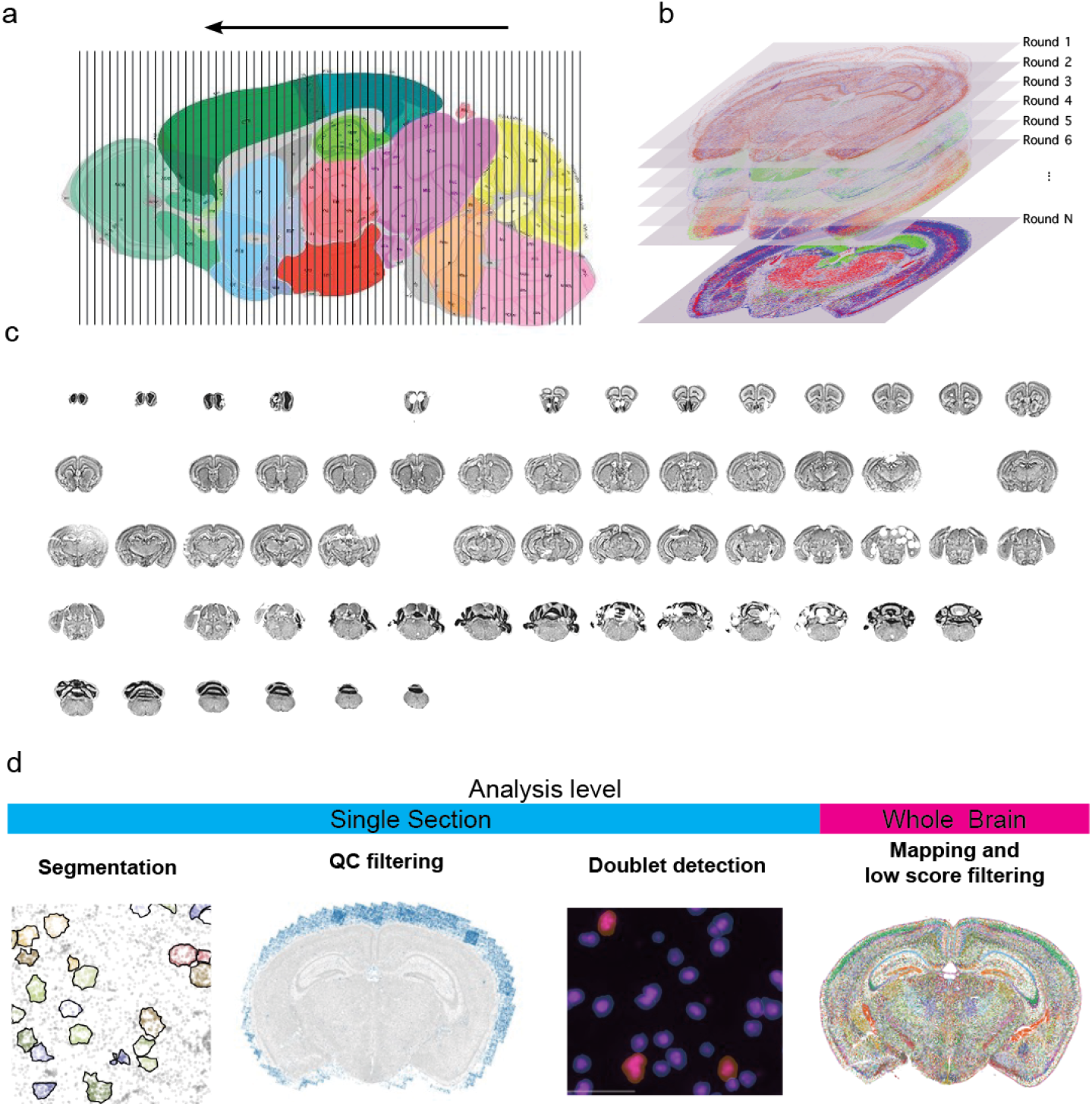
Overview of data generation and processing for the ABC atlas data. (**a**) Sampling plan for whole mouse brain dataset. The brain was sectioned from posterior to anterior and sections were collected ever 200 µm. (**b**) For each section 500 genes were imaged. (**c**) Sections after initial quality control using MERQUACO (Martin et al., 2025). Gaps indicate sections that failed. (**d**) Data processing steps in order. While the first 3 steps can be performed at the individual section level, the final mapping and low-quality filtering are performed on the whole dataset.

Beyond the automated steps, we provide additional code modules to enable more advanced analyses. These include tools for subsetting the dataset into spatially coherent and anatomically meaningful regions using spatial domain detection, as well as registration of the data to the mouse brain common coordinate framework version 3 (CCFv3) via landmarks based on anatomically restricted cell types. Finally, we offer gene imputation functionality using the ENVI package (Haviv et al., 2025), which can be run as an optional extension to further enrich the dataset. This modular approach allows users to benefit from a robust automated pipeline for core processing, while also giving flexibility to perform custom or advanced analyses using the provided additional code.

### Segmentation

Cell segmentation assigns individual transcripts to cells. Segmentation involves delineating the boundaries of each cell within a tissue image, which can be challenging due to the complex and heterogeneous nature of tissue structures. In Yao et al. (2023b), we segmented cells for our dataset using the VizgenPost-processingTool (VPT, https://github.com/Vizgen/vizgen-postprocessing) and Cellpose segmentation family with the cyto2 model properties (Stringer et al., 2021). The cyto2 model has two input channels, DAPI for the cell body and PolyT for the cytosol (Fig. 2e). This method has several shortcomings. 1) Segmentation was performed on a single plane that had to be manually defined (generally, we used the central plane of the image stack) and cell outlines were applied to the planes above and beyond leading to a cylindrical cell shape (Fig. 2a). 2) Overestimation of cell size, exemplified by low transcript density areas in the periphery, could cause contamination with extracellular mRNA (arrows in Fig. 2a). 3) Erroneous detection of cells outside the brain, likely due to detection of out-of-focus beads in the PolyT channel (arrows in Fig. 2c). 4) Under-segmentation in high density areas such as the cerebellum (red boxes in Fig. 2c). 5) Inaccurate somatic outlines with PolyT (Fig. 2e). To improve cell segmentation we updated the cyto2 model with MERSCOPE data from various species (mouse, macaque, and human) using the human-in-the-loop approach introduced with Cellpose2.0 (Pachitariu and Stringer, 2022), created a software package to run Cellpose segmentation in 3D (Spots-in-Space, SIS, see methods) and replaced PolyT images with images of total mRNA density (Fig. 2e). Using this approach, we saw a significant improvement in cell segmentation quality. First, visual inspection revealed cell outlines that matched our expectation of somatic shapes in brain tissue, without “padding” around the cells (Fig. 2b, arrows). Second, improved detection of small somata in regions of high cell density such as the granule cell layer in cerebellum (red box, Fig. 2d). In some regions, VPT segmentation detected no somata where our Cellpose model detected many (black arrows, Fig 2c). Third, reduced detection of off-tissue cells. As expected, the transition from cylindrical VPT to 3D Cellpose segmentation reduced genes per soma (from 103 to 84, Fig. 2f), transcripts per soma (from 332 to 257, Fig. 2g), and cell volume (median reduced from 1037.54 µm3 vs 473.06 µm3, Fig. 2h), while within-soma transcript density is increased (median increased from 0.35 to 0.81 transcripts/µm, Fig. 2i). The increase in transcript density indicates that the regions of the cell removed are primarily low-density regions at the periphery of the cell. To determine whether the revised segmentation resulted in a cleaner gene profile, we looked at the expression of incongruent genes, genes whose expression is mutually exclusive within cells. By subsetting the single-cell RNAseq taxonomy data to genes present in the MERSCOPE panel, we identified marker genes at the division level using a standard differential gene expression workflow (see methods, Fig. S1a), and then created a list of incongruent gene pairs by combining markers from different members of the division (Table S2). For each cell we calculated the presence of incongruent genes as a percentage of total detected transcripts. 3D Cellpose segmentation reduced incongruent genes relative to VPT segmentation (Fig. 2j). The difference is more pronounced when comparing only highquality cells (Fig. S1b). A prominent example is the reduction in transcripts of the vesicular glutamate transporter Slc17a7 in GABAergic cell classes CTX-MGE GABA and CTX-MGE GABA (Fig. S1c), which is expected since Slc17a7 is in processes that traverse the neuropil (Glock et al., 2021). Similarly, improved segmentation reduced the incidence of GABAergic markers in glutamatergic cells (Slc32a1 and Gad2 in IT-ET Glut and NP-CT-L6b). Hence, 3D Cellpose segmentation resulted in cleaner gene expression profiles and increased correlation scores during label transfer (Fig. 2k).

**Figure 2.**
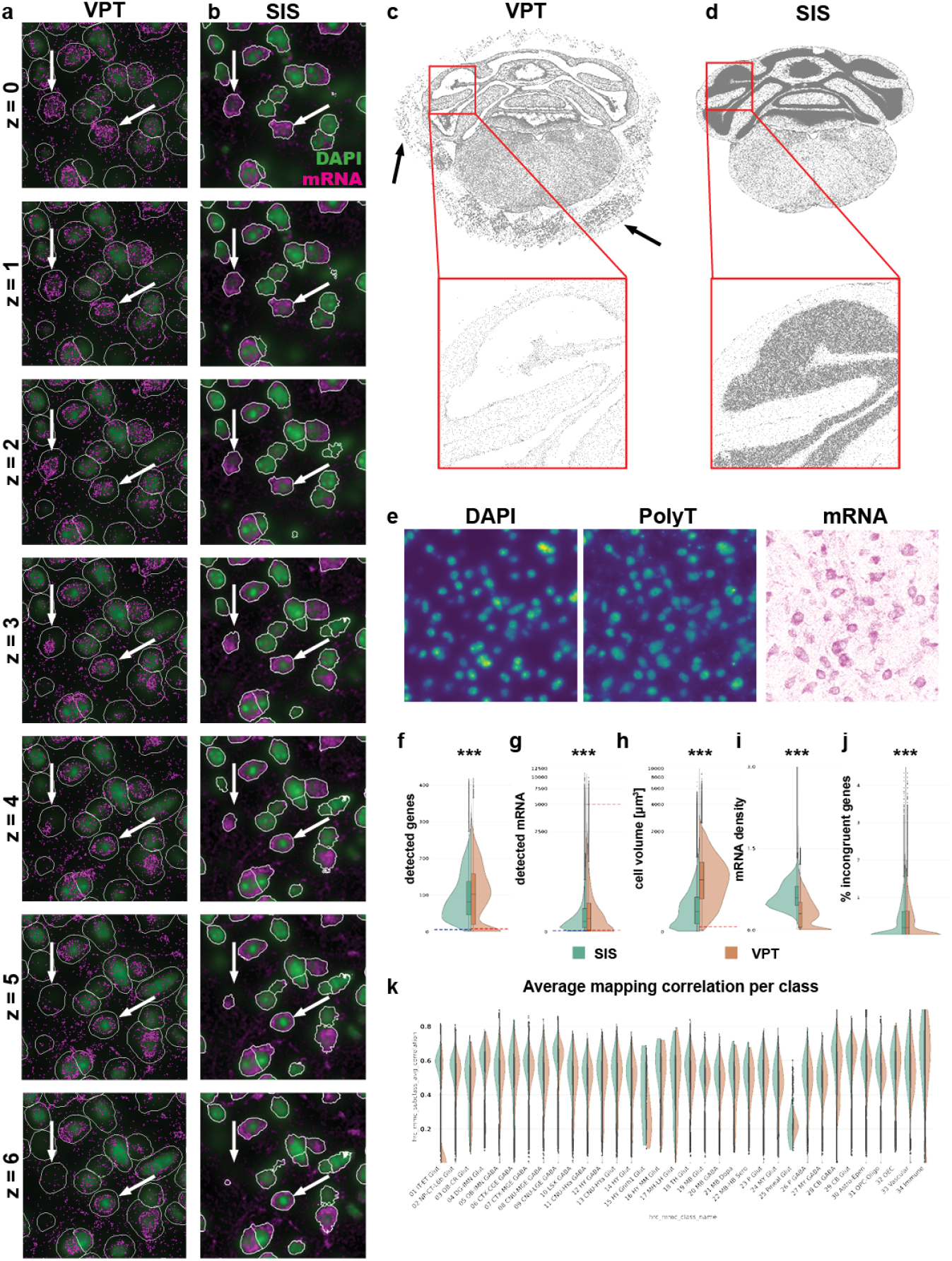
Cell segmentation. (**a** and **b**) Example segmentation from Vizgen Post-processing Tool (VPT, a) and Spots in Space (SIS, b). Images show the same region for comparison. Cell outlines are in white. Arrows indicate examples where SIS results in more realistic 3D segmentation. Each image represents a different imaging plane. (**c** and **d**) Example demonstrating improved detection of cells in dense area for SIS (c) compared to VPT (d). (**e**) Images showing staining for DAPI and PolyT as well as density image generated from detected mRNA. The mRNA image highlights that PolyT is not faithfully recapitulating the entire extent of a cell cytosol. (**f**-**j**) Violin plots of main cell quality criteria for all cells in the ABC-atlas dataset segmented either by SIS (green shade) or VPT (orange shade). The values for genes (f) or transcripts (g) per cell, volume (h) and incongruent genes (j) are significantly decreased, the mRNA density (i) per cell is lower. All these are indications for cell of higher quality (see text for explanation) (**k**) Distribution of correlation coefficients within class assignment (explained later in the manuscript). ***P < 0.001, Two-sample Kolmogorov-Smimov-Test

Cell volume serves as a critical parameter for distinguishing cell types, and our enhanced segmentation approach yields cell volumes that more accurately reflect biological reality. To assess the added value of these improved measurements, we examined cell volume distributions by leveraging a recently published electron microscopy (EM) dataset of mouse visual cortex, which provides subclass-specific cell volume estimates (Elabbady et al., 2025). Given that cell boundaries in our study are defined using total mRNA images—and the gene panel is highly enriched for neuronal markers, we focused our analysis on neuronal subtypes. Notably, we observed that excitatory layer 5 extratelencephalic (5P-ET) cells exhibit larger volumes compared to other subclasses, whereas excitatory layer 4 (4P), layer 5 near-projecting (5P-NP), and inhibitory bipolar cells (BPC) display smaller volumes (Fig. S2a). These particular subtypes were selected for comparison as they represent the extremes of the volume distribution. We identified the corresponding subclasses between the EM dataset and our taxonomy (Table S3). When comparing the larger subclass to the smaller ones, we observed distinct differences in volume distributions that became even more pronounced with SIS segmentation. The median cell volume was reduced in SIS compared to VPT segmentation (Table S4). However, the differences between subclasses in volume distribution were more marked with SIS (as quantified by Cohen’s D, Table S5), highlighting improved subclass separation (Fig. S2b-d).

### Low-quality filtering

The improved segmentation reduced the number of lowquality cells, although additional factors can still produce such cells. For example, somata cut at the section surface contain only part of the soma, resulting in partial transcriptome measurements. Remaining low-quality cells were removed using thresholds of six genes per soma and 30 transcripts per soma (Fig. 2f and g, blue dashed line). The thresholds were manually selected to fall within the outlier portion of the distribution of each feature. Plotting the spatial distribution of low-quality cells revealed that most were in regions of poor tissue quality, particularly where tissue detached from the coverslip (arrows in Fig. S3a and b), further validating the choosen cutoffs. Compared to the previous segmentation, the new segmentation yielded lower thresholds (genes < 15, transcripts < 40; Fig. 2f and g, dashed red line on the right side of the split violin plot), presumably because (1) fewer false negatives (off tissue cells) occur, and (2) more restrictive cell boundaries reduce contamination from extracellular or neighboring transcripts, which often have distinct gene profiles and artificially increase the number of unique genes per cell. In the original dataset, cells below a certain volume (< 100 µm^3^; Fig. 2f) were also excluded. However, the volume distribution obtained with the new segmentation showed no clear lower threshold, so this filter was not applied. We previously removed cells with unusually high transcript counts, assuming they represented doublets (Fig. 2g), but replaced this approach with a dedicated doublet detection method (see below). We also identified a third indicator of low-quality cells based on the percentage of blank barcodes. Blank barcodes correspond to entries in the MERSCOPE codebook that satisfy all codeword criteria (same Hamming distance and weight) but do not map to targeted genes, serving as a measure of falsepositive detection. The distribution of blank counts per cell showed that most cells contained fewer than 2% blank barcodes (Fig. S3d, red dashed line), and this pattern was independent of the segmentation method (Fig. S3d). Regions with poor tissue quality, such as detached areas, exhibited elevated blank barcode levels (Fig. S3c). After applying all filters, we removed approximately 600,000 cells, representing less than 10% of the original dataset of 6.5 million cells.

### Doublet removal

Doublets occur when two or more cells are in proximity and are identified as a single cell. To identify doublets in our dataset, we used the deep-learning based method Solo (Bernstein et al., 2020). Briefly, Solo generates artificial doublets. Expression profiles including the artificial doublets are placed in a latent space using a variational autoencoder (VAE) and a classifier is trained on the embedded data to distinguish singlets and doublets. The classifier is applied to the original data embedded in latent space, and the results are singlet and doublet probabilities for each cell. We calculate the difference (dif) between singlet and doublet scores and determine the thresholds for doublet as quantile 0.9(dif) – quantile 0.1(dif) (Fig. 3a, red dashed line). Visual inspection revealed that many doublet cells contained multiple nuclei (Fig. 3b, red boundaries). Doublets also contained more genes (Fig. 3d) and transcripts (Fig. 3e), were of greater volume (Fig. 3f), and contained more incongruent genes (Fig. 3g). Typically, doublets were evenly distributed across a section (Fig. 3c) at low density, with only 180,000 doublets detected in a dataset of 5.9 million cells.

**Figure 3.**
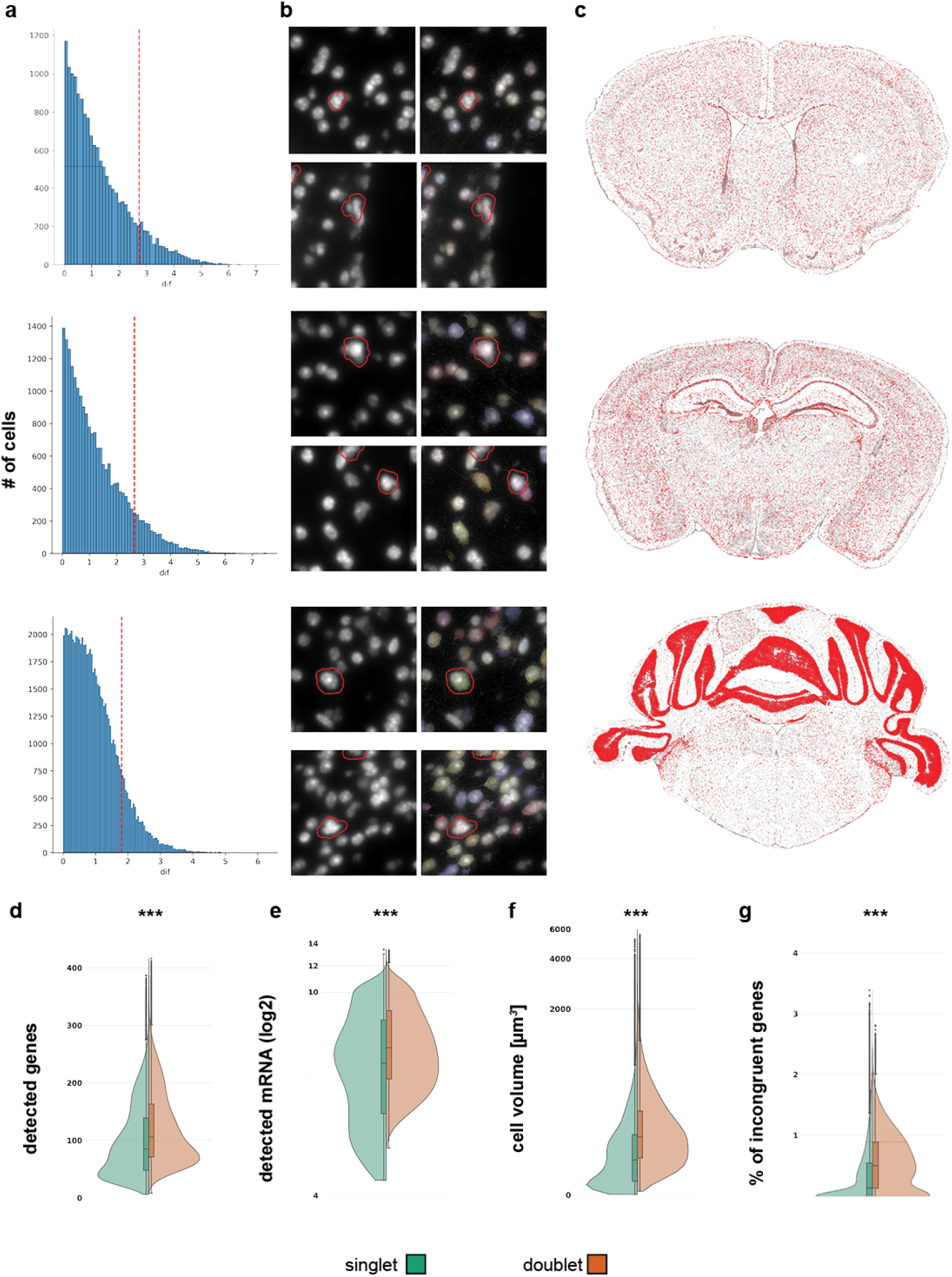
Doublet removal. (**a**) Histogram for the distribution of difference between singlet and doublet score. Red dashed line indicates cutoff for each section. (**b**) Example DAPI images of zoomed in regions. Red outlines are highlighting cells that are classified as doublets. (**c**) Overview of example sections to illustrate spatial distribution of doublets. Cells identified as doublet are shown in red. No obvious hotspots are discernible. (**d**-**e**) Comparison of distribution for cell quality metrics in cell identified as singlets (green hue) and doublets (orange hue). Doublets consistently had statistically higher values for genes (d), mRNA (e) spots, cell volume (f), and incongruent genes (g).***P < 0.001, Two-sample Kolmogorov-Smimov-Test

### Label transfer

We performed label transfer from the Allen adult mouse brain taxonomy (Yao et al., 2023b) to cells in our MERSCOPE dataset using MapMyCells (RRID:SCR_024672), which provides a correlation score that indicated the robustness of each label. In Yao et al. (2023b), we applied a threshold of 0.5 to cluster labels to remove low quality cells. However, the median and distribution of average correlation coefficients can differ substantially between clusters (Fig. 4a), which is caused by the variable number of cell type specific markers for individual clusters. As a result, a fixed threshold might exclude some accurately mapped cells and retain low quality cells of a different cluster. One example is supertype 0682 RN Spp, the majority of which were eliminated by the fixed threshold (red cell in the left panel of Fig. 4b). To identify outliers with low average correlation coefficient in a more principled way, we used a specialization of the median absolute deviation (MAD), the doubleMAD, a measure that is often robust for skewed distributions. We calculated upper and lower MADs (MAD^high^ and MAD^low^) based on deviation of values above or below the median of the distribution. We removed cells with an average correlation value lower than the median minus 3*MAD^low^ (red dashed lines, Fig. 4a). In the case of the supertype 0682 RN Spp, this revised approach retained 785 of 831 cells (green cells in the right panel of Fig. 4b).

**Figure 4.**
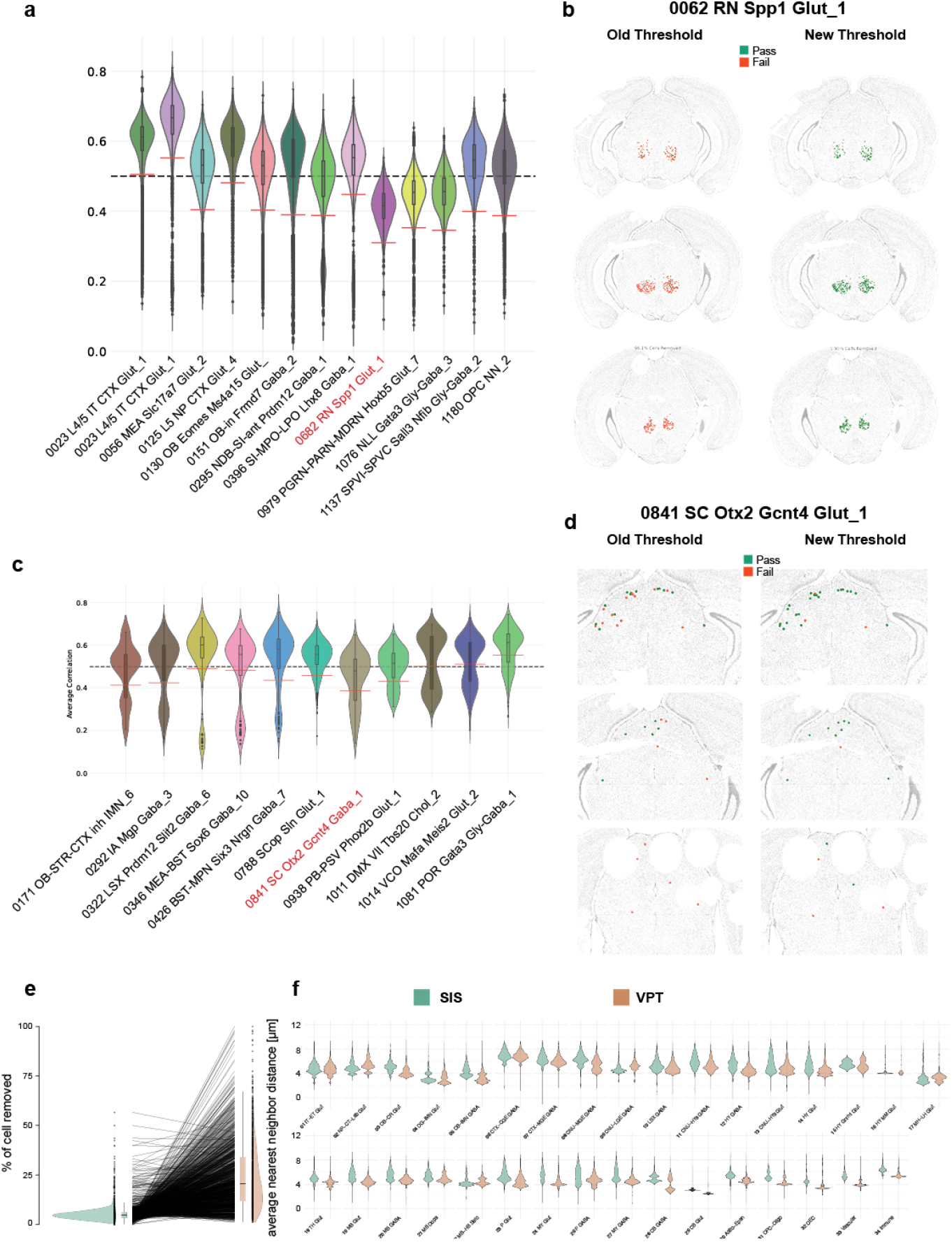
Filtering of cells with low cell type assignment confidence. (**a**) Example distribution of correlation coefficients for selected supertypes. Black dashed line indicates the original fixed threshold of 0.5. Red solid lines indicate new doubleMAD based supertype specific threshold. (**b**) Example of retained cells from supertype highlighted in red in a. Left-hand side is the old threshold right-hand side the new doubleMAD threshold. Cells in red are filtered out and cells in green are retained. (**c**) Example distribution of correlation coefficients for selected supertypes with bimodal distribution. Red dashed lines indicate new doubleMAD based supertype specific threshold. (**d**) Example of retained cells from supertype highlighted in red in c. Left-hand side is the old threshold right-hand side the new doubleMAD threshold. Cells in red are filtered out, and cells in green are retained. (**e**) Failure rate per supertype as measured by percentage of cells removed by either fixed threshold or adaptive doubleMAD threshold. (**f**) Distribution of average nearest neighbor distance within supertypes of filtered cells grouped by class.

More complex situations arise when distributions display a bimodal shape (Fig. 4c). Potential causes could be a higher tendency of cell types to be confused with low-quality cells or alternatively, we are mapping to a separate cell type not present in the scRNAseq taxonomy. To overcome this problem, we determined the local minimum between both peaks and discarded the lower distribution (see Methods). For the remaining distribution, we applied the procedure described above. For example, supertype 0841 SC Otx2 Gctn4 Gaba_1 displayed two peaks in the average correlation coefficient distribution, the second driven by low quality cells (Fig. 4c). The original threshold of 0.5 would remove those low-quality cells and cells within the superior colliculus (SC, red cells left side Fig 4d). The new filtering method removed cells in low tissue quality regions and retained a greater proportion (122/173 vs 71/173) of cells in the SC. Overall, of the 1201 supertypes, more cells were retained in 999, fewer in 192, and no change in 10 (Fig. 4e, Table S4). We previously demonstrated that cell types have a high degree of regional localization (Yao et al., 2023b). We reasoned that poorly mapped cells lack this feature. To examine this, we quantified the average-neareast-neigbbor (ann) distance within supertypes for each cell removed by either method. When we plotted the distribution of all ann distances split by class we could observe that in a majority of cases the cells removed by the MAD-based threshold have higher values (Fig. 4f). Overall, this method results in a higher retention of cells per supertype than the fixed threshold method. The maximum observed increase in positive retention is 92.6% (0682 RN Spp), while the largest reduction in the number of cells per supertype is 13.9% (0334 LSX Sall3 Lmo1 Gaba_4). Of the top 10 supertypes with more cells retained, 6 belonged to non-neuronal classes (Table S6). With fewer non-neuronal markers in our gene panel, we typically detect fewer transcripts in non-neuronal than neuronal cells, making non-neuronal cells more susceptible to low correlation scores. Using this method, we end up removing an additional 300.000 cells leaving us with 5.5 million high quality cells. Alternatively, when applying the old, fixed threshold we would have removed 1 million additional cells.

### Spatial domain detection

Analyzing data at the whole-brain level presents several challenges. Brain regions differ in features such as cell density, neuron-to-glia ratio, and excitatory–inhibitory balance, which complicates the interpretation of metrics like cell–cell interactions, neighborhood enrichment, or ligand–receptor relationships. To address these issues, identifying spatial domains is a crucial step. Spatial domain detection involves recognizing regions that are spatially contiguous and share similar molecular characteristics, such as gene expression profiles. This enables a more biologically meaningful analysis by grouping cells into functionally and molecularly coherent units. Machine-learning (ML) tools based on graph-neural networks can efficiently capture local spatial and molecular relationships. These ML tools generate a latent embedding of spatial transcriptomics data (spots or cells) based on the gene expression of itself and its immediate neighbors. The resulting embedding can be utilized by standard clustering algorithms (e.g., Louvain or Leiden) to generate computationally derived spatial domains. We used STAligner, a graph attention neural network designed for integrating and aligning spatial transcriptomics datasets (Zhou and Zhang, 2023). STAligner accepts a stack of sections as input and finds spatial neighborhoods extending across multiple sections. We input transcript locations, enabling STAligner to incorporate extra-somatic transcripts such as those in axons, dendrites, and white matter tracts, which can be helpful in discriminating anatomical regions. We first created a spatial neighbor graph for each section. To reduce computational cost, the transcript table was downsampled to 30 µm grids (Fig. 5a, left). Grids were filtered based on quality criteria similar to the ones used for cells (Fig. S4a, see Methods). A latent embedding of cells was generated using STAligner with the neighborhood graph set to eight nearest neighbors (Fig. 5a, middle and right). Leiden clustering was performed across resolutions from 0.2 to 2.0 in 0.1 increments, and cluster stability, measured by Adjusted Mutual Information and Adjusted Rand Index, remained consistent (Fig. S4ab and c). Resolutions between 0.8 and 1.6 were inspected further, and spatial domains were transferred to the cell-by-gene table. We selected a resolution of 1.4 as it best separated neuronal from non-neuronal regions, providing a balance between anatomical detail and interpretability. We identified 67 initial spatial domains (Fig. 5b and c, Table S7). 4 were identified as low-quality, either because they co-occurred with lowquality tissue regions (as exemplified by the lower genes per cell, 70, and transcripts per cell, 196, Fig. S4d-f) or because they contained very few cells that were dispersed across the brain and were removed. Two other domains (Borders_1 and Borders_2) were located at the periphery of the brain, were dominated by vascular cell subclasses (Fig. S4g and h), and removed before further analysis. At this resolution most of the fiber tract spatial domains are specific to white matter. However, we still find regions that contain neuronal cell types. For example, spatial domain fiber_tracts_3 contained not only white matter regions, such as the internal capsule, but also the reticular nucleus of the thalamus (RT) (Fig. S4i and j). The UMAP shows a high degree of intermixing among sections (Fig. S5a), demonstrating that the latent embedding captures the gradual change in regional composition from anterior to posterior. The main exception is the anterior group of sections (orange hues), which are dominated by olfactory areas (OLF). This indicates that no batch effect was introduced by sectioning, as most regions are represented across multiple sections and transition smoothly along the anterior–posterior axis. The anatomical localization of spatial domains ranged from broad regions (e.g., P-MY_1, Fig. S5b) to highly specific areas (e.g., PSV_1, IC_1, Fig. S5c and d). These domains are characterized by distinct subsets of subclasses (Fig. 5d), with posterior regions (MB-P and P-MY) containing a greater diversity of subclasses compared to more anterior regions (Isocortex-CLA, HIP). To validate our method, we primarily assessed whether the spatial domains corresponded to known anatomical regions based on regionspecific cell types. Our main objective was to confirm that established anatomical landmarks, defined by distinct cell-type compositions, were accurately represented within individual spatial domains and did not cross domain boundaries. To quantify the concurrence of spatial domains with known anatomical regions, we divided the brain into 18 main regions and identified cell types specific to these regions (Fig. S6a, Table S8). We calculated the overlap of cell types between spatial domains and assigned brain regions using the Jaccard coefficient and found that most spatial domains were primarily assigned to 1 or 2 broad regions (Fig. 5e). This approach increased our confidence in the reliability of the spatial domains as meaningful anatomical units. We also noted that regions sharing similar cell-type profiles were often merged into single domains. For instance, some spatial domains (OLF–ENT–EP) included cell types from multiple areas, such as the olfactory (OLF), entorhinal (ENT), and endopiriform (EP) regions, reflecting their shared cellular characteristics. In contrast, areas like the hippocampus (HIP) and thalamus (TH) were well-confined within distinct spatial domains. Following these validations, we aggregated the spatial domains into 19 broader regions, ensuring that most aggregated domains remained within specific anatomical regions, with only a few spanning two neighboring areas—most notably in the posterior brain. This aggregation provided a consistent and interpretable framework for subdividing the whole-brain dataset (Fig. 5f and g)

**Figure 5.**
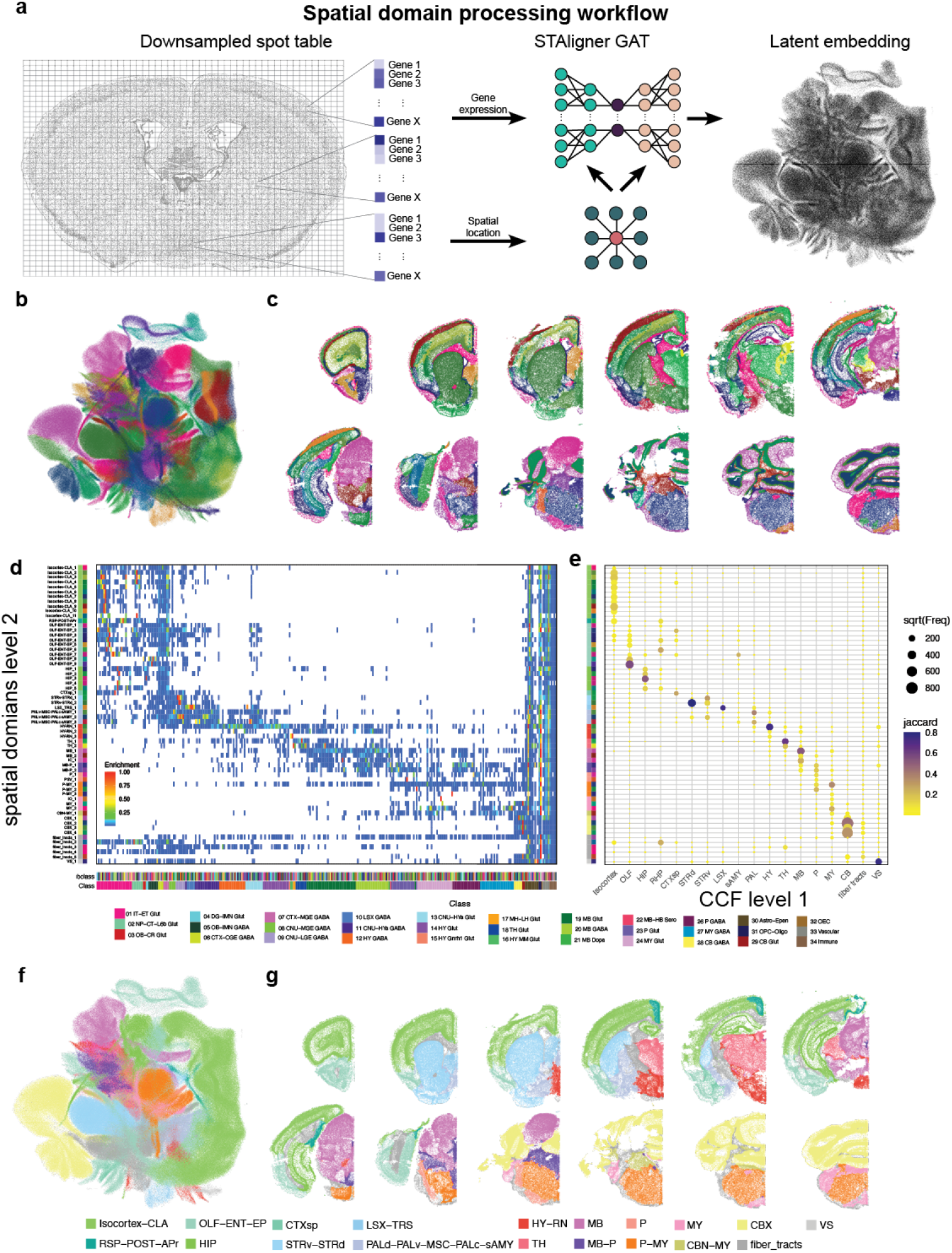
Spatial domain detection. (**a**) Workflow for creating latent embedding using STAligner. (**b**) UMAP of STAligner latent embedding colored by spatial clusters obtained from Leiden clustering. (**c**) Example sections labeled by spatial clusters. For space reasons, only one hemisphere of the section is shown. (**d**) Heatmap showing subclass enrichment in spatial domains. (**e**) Dotplot demonstrating the Jaccard overlap of region-specific cell types in spatial domains. Dot size indicates number of cells overlapping and color indicated Jaccard coefficient (f) UMAP colored by higher level of spatial domains aggregated by dominant CCF region per domain. g) Example sections labeled by higher level of spatial domains.

### Registration

While assigning cells to spatial domains is helpful for downstream analysis, to integrate spatial transcriptomics data successfully with other modalities, such as single-neuron morphologies (Peng et al., 2021), Patchseq (Gouwens et al., 2020; Sorensen et al., 2023), and viral tracer studies (Oh et al., 2014; Yao et al., 2023a), the data needs to be registered to the Allen Mouse Brain CCFv3. A key requisite for good registration is a reference channel with high information content. Classically, registration is performed using either background autofluorescence or specifically selected fluorescent markers. In the case of our MERSCOPE experiments, the only available reference signals are DAPI and PolyT. Both signals lack texture information that adequately reflects the anatomical complexity of the mouse brain (Fig. S7a and b). To overcome this, we decided to register not using a reference stain but instead use the region annotation of the Allen Reference Atlas (ARA). For this we made use of the broad region assignment described earlier, converting the cell centroids into images to be used as reference stain. To register the data to the CCF, we used QuickNII and VisuAlign for manual alignment based on CCF region annotations (Puchades et al., 2019, 2025). In addition to broad region assignments (see above), we further identified clusters marking 53 additional landmark regions (Fig. S7c, Table S9). Images were generated for each section to visualize cell assignments (Fig. 6a), and section orientations were adjusted to account for mounting variability (see methods). We then applied both linear (using QuickNII, Fig. 6b) and non-linear transformations (using VisuAlign, Fig. 6c) to compensate for section tilt, tissue deformation and individual differences, enabling accurate plotting of cell centroids within our 3D reference space (Fig. S7d). When we plot representative cell types with a high degree of spatial localization in our 3D reference space, we can see those cell types are restricted to the region they are associated with (Fig. 6d). Next, we plotted the enrichment of subclasses per region as a heatmap. We focused on neuronal classes and grey matter and disregarded non-neuronal cells or white matter regions. Our goal was to optimize the registration of neuronal cells to grey matter. Very similar to our initial result (Yao et al., 2023b) we observe a high spatial correlation of transcriptomically similar cell types, with groups of classes being restricted to broad anatomical region (Fig. 6e). To assess registration improvements, we applied the original Yao et al. (2023b) registration to the newly processed dataset. The updated registration yielded 660,052 additional cells not present in the previous registered dataset, while 371,120 cells from the original dataset were absent in the new registration, resulting in a net gain of 288,932 cells. Most newly added cells were due to the successful registration of four additional sections (Fig. S8a), with remaining gains primarily in areas bordering white matter regions, reflecting improved registration accuracy. Quantification revealed that the majority of newly registered cells are neurons, particularly cerebellar neurons and immature neurons from the olfactory bulb (Fig. 6g). Comparison of cellular enrichment between registrations showed no substantial differences, with minor changes restricted to broader anatomical regions (Fig. 6f). To assess the accuracy of the registration, we calculated the overlap of regionspecific clusters within their predicted regions (Fig. 6h). The overlap exhibited a high degree of accuracy, as indicated by the Adjusted Rand Index (ARI) of 0.97. This analysis was also performed for cell types specific to smaller regions (Fig. 6i), demonstrating a high concordance between the expected locations of cells and their registered regions in the CCFv3 (ARI = 0.95). A similar analysis was conducted on the dataset with the previous registration (Fig. S8d and e). The differences were minimal, with the new registration showing a slight improvement at the broad region level (0.97 vs 0.96), while no difference was observed at the smaller region level (both ARI = 0.95). Given that registration errors were minimal during quantification, we conducted a detailed analysis of instances of misalignment. In one instance in the hindbrain, the initial registration included a section where cells of the inferior colliculus (IC) were incorrectly assigned to the cerebellum (CB). Manual adjustment of section tilt resolved this issue (Fig. S8b and c). In summary, while quantitative assessments of registration accuracy are useful, they have limitations because visually apparent but small errors may go undetected. Therefore, visual inspection remains essential for identifying residual registration issues that quantitative measures might miss.

**Figure 6.**
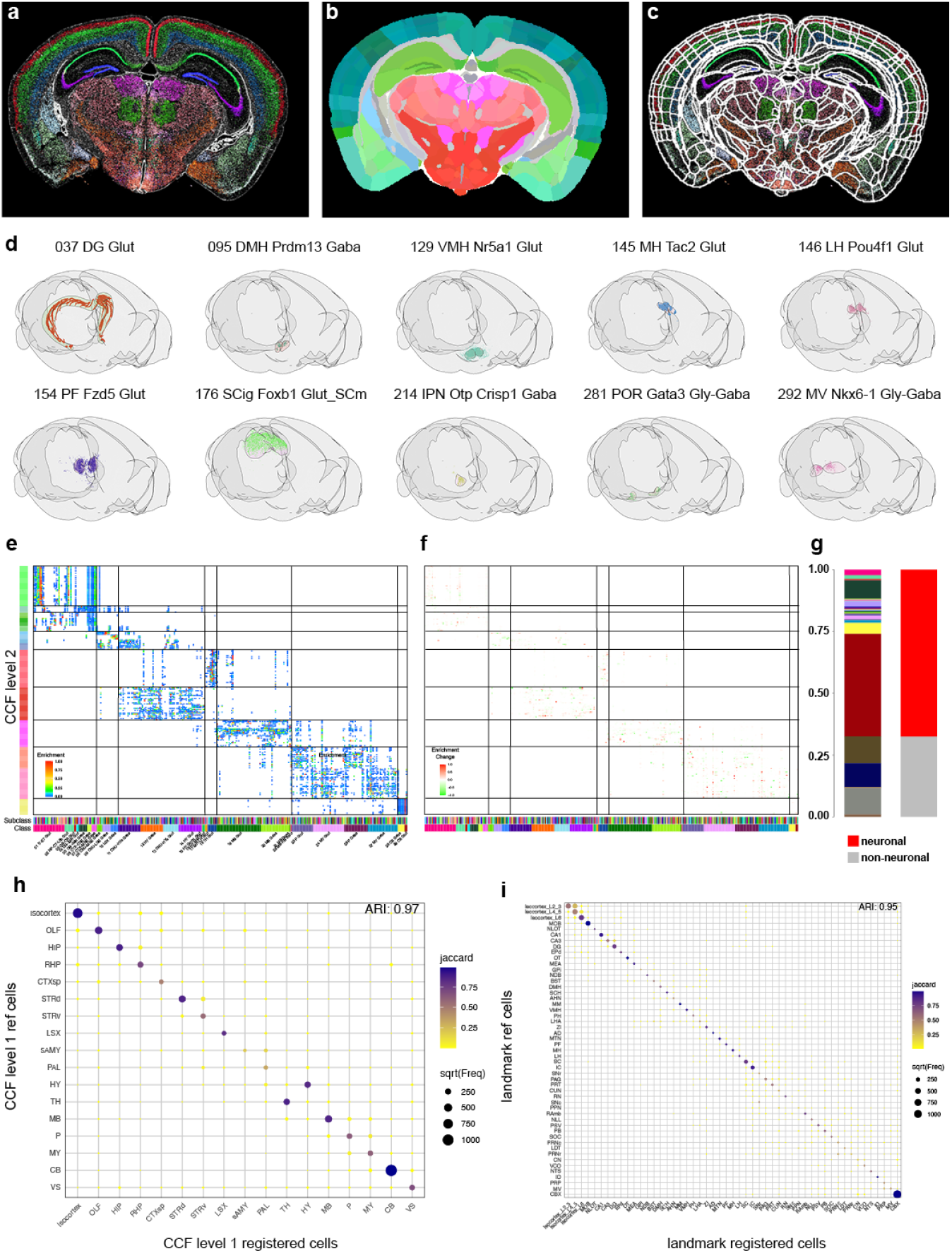
Registration of MERSCOPE data to CCFv3. (**a**) Example image of cell centroid locations. Cells are colored based on regions they are specific to. (**b**) Matching CCFv3 section adjusted for tilt and scale using QuickNII. (**c**) Boundary overlay of CCFv3 regions after non-rigid landmark-based adjustment using VisuAlign. (**d**) Representative example of registration accuracy of region-specific subclasses in final 3D CCFv3. (**e**) Heatmap showing enrichment of subclasses within individual CCF regions. (**f**) Change in enrichment score compared to registration used in Yao et al. (2023b). (**g**) Stacked bar chart showing the distribution of added cells colored by class (left) or neuronal vs. non-neuronal (right). (**h**) Dotplot highlighting the overlap of broad specific cells and their CCF region assignment after registration. Size indicated number of cells and color Jaccard coefficient. ARI = Adjusted Rand Index. (**i**) Same as in h but instead of broad regions more fine grain landmarks are compared.

### Gene imputation

While it is possible to map expression of the entire ∼32,000 genes in the mouse genome using imaging-based spatial transcriptomics techniques (Cohen et al., 2025), the vast majority of datasets include only hundreds of genes. Our dataset imaged 500 genes, and we performed imputation to estimate the likely locations of additional genes. We applied gene imputation to all the sections of the dataset using the COVET/ENVI framework (Haviv et al., 2025). Gene imputation was performed separately for each of the 18 major brain regions using cells from relevant sections. Matching scRNAseq cells were selected for each region and subsampled. Both spatial and scRNAseq datasets were coembedded, and an ENVI decoder was used to impute missing gene expression (Fig. 7a, see methods). To illustrate the successful imputation of gene expression we compared the imputed gene expression to the originally measured gene expression and a representative section from the Allen ISH atlas (Lein et al., 2007) (Fig. 7b, for genes present in the panel) or only to the Allen ISH atlas (Fig. 7c. for genes not present in the panel). To quantify imputation we made use of an unpublished whole brain dataset that was generated with a different gene panel containing 263 genes not present in our current panel. We identified matching sections between the two datasets (see methods) and used CAST (Tian et al., 2023) to have both sections in the same anatomical space (Fig. S9a). We calculated the Multiscale Spectral Similarity Index (MSSI) from the ENVI package. The MSSI assesses the spatial agreement of gene expression patterns by incorporating how close cells are to each other and analyzing the pattern at various scales. When we plotted select genes with a high MSSI index we found again high concordance of imputed gene expression and measured gene expression (Fig. S9b). When we plotted the distribution of MSSI values for all 263 genes either split by hemisection (Fig. S9c) or a mean for the whole dataset (Fig. S9d) we found that most genes have a MSSI above a value of 0.75. The added genome-wide coverage will allow us to interrogate the spatial expression of functionally relevant genes (ion-channels, GPCR’s, cell-adhesion molecules, etc) which are often underrepresented in gene panels to accommodate cell type markers.

**Figure 7.**
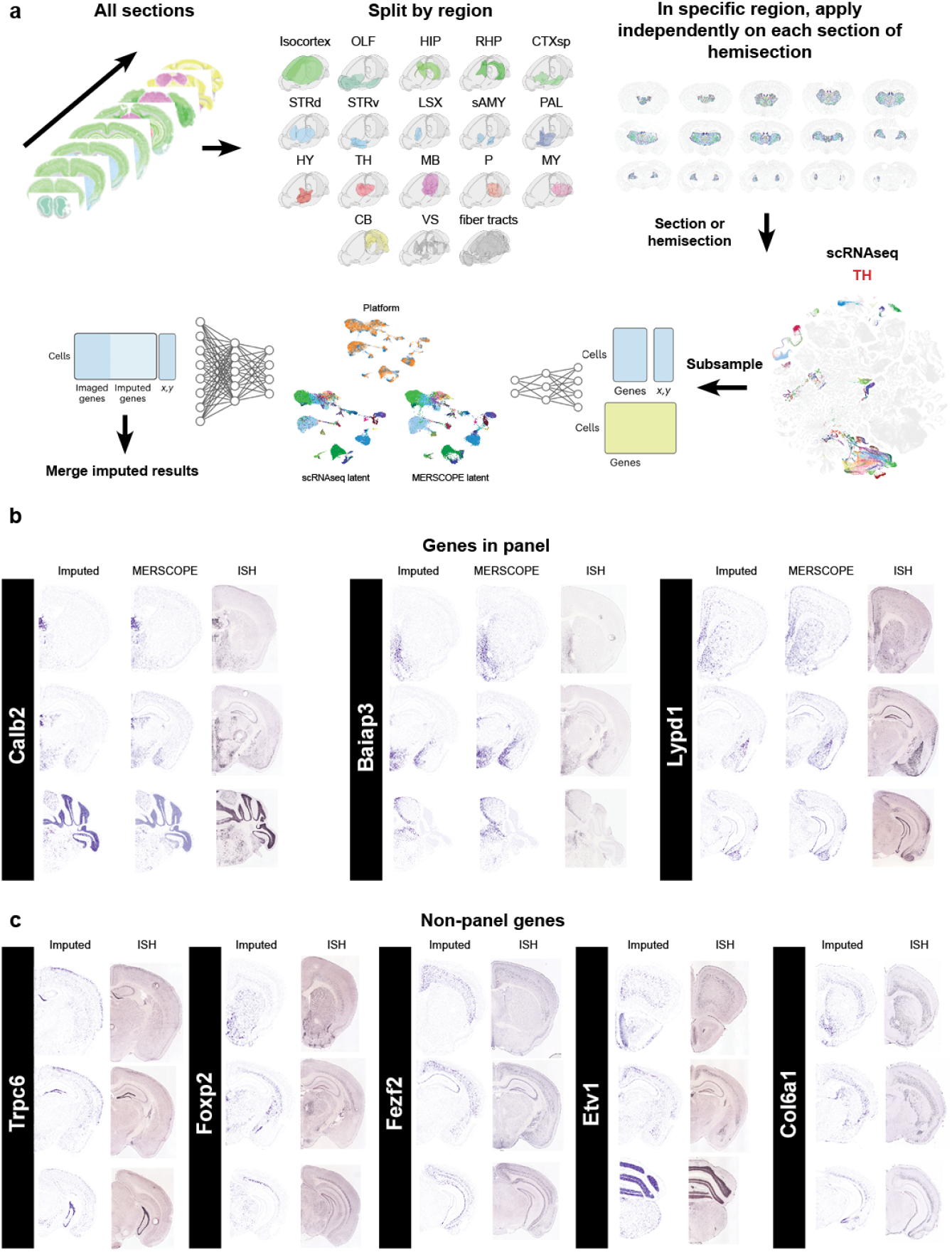
Gene imputation on MERSCOPE data. **(a)** Workflow for imputing genome-wide mRNA expression from scRNAseq data. We use all sections and split them up by main regions. To reduce computational load, we selected cells from one section or hemisection from each region and integrated it with subsampled scRNAseq data from the matching region. Data gets transformed into combined latent space using the encoder. Missing gene expression is imputed from scRNAseq data onto the MERSCOPE data using the decoder. **(b)** Examples of imputed gene expression for genes present in the panel showing imputed expression (left), original MERSCOPE expression (middle) and comparative section from the ISH atlas (right). (**c**) Example of imputed gene expression for genes not present in the panel showing imputed expression (left) and comparative section from the ISH atlas (right).

### Applicability to other methods

Our automated pipeline is readily adaptable to datasets generated from alternative platforms such as StereoSeq and Xenium, as well as other tissue types including human cortex samples and developing mouse brain (Fig. S10). In each application, it is necessary to adjust the threshold parameters to account for platform-specific data characteristics and employ specialized Cellpose models tailored to the imaging modality and tissue type. The output formats produced by our pipeline are compatible with the other analysis code described in this manuscript, except for the registration step, which requires the manual identification of cell-type landmarks and the existence of a CCF and standard brain.

## Discussion

We present an optimized and streamlined processing pipeline for cellular resolution spatial transcriptomics data analysis. This workflow encompasses cell segmentation, quality control filtering, doublet detection, cell type mapping, spatial domain analysis, and registration to the Allen Mouse CCFv3. Each module is designed to be flexible and can be adapted for use with other platforms that offer cell or subcellular resolution. Enhanced cell segmentation enables more precise identification of individual cells within complex tissues, which is essential for the accuracy of downstream analyses. Quality control thresholds are empirically established and iteratively refined for each dataset to adjust for platform specific detection efficiency, ensuring that only high-quality data are included in the final analysis. To validate the effectiveness of our new pipeline, we applied it to the existing MERSCOPE ABC-WMB dataset and demonstrated measurable improvements in cell registration accuracy and overall data quality. A key improvement in our pipeline is enhanced cell segmentation, which produces more accurate cell boundaries and cleaner gene expression profiles by using mRNA images to mark the cytosol. However, because the current gene panel is biased toward neuronal genes, immune cells are difficult to detect using this approach and may be underrepresented. Expanding future gene panels to include markers for these cell types will improve their identification and address this limitation. The implementation of our new segmentation approach has revealed more pronounced differences in cell volumes, offering enhanced resolution in distinguishing individual cellular structures. However, it is important to note that total measured cell volumes in MERFISH datasets are likely attenuated. This attenuation arises because the majority of cells in our samples exceed the 10 µm section thickness typically used in MERSCOPE imaging, resulting in limited cell profiling and potential underestimation of actual cellular dimensions. We are presenting the filtering criteria employed in our analysis, along with the rationale for their application. These criteria are not static; rather, they can be adjusted on a project-specific basis and must account for the particular platform, gene panel, and tissue type utilized in each study. We employed spatial domain detection software to subdivide our data into biologically meaningful subsets. The software allowed us to identify clusters of cells with similar gene expression profiles and anatomical locations. To ensure the biological validity of these subsets, we performed rigorous verification against known anatomical markers, confirming that our method accurately reflects the complex organization of the tissue. The resulting data can be leveraged for more region-specific, in-depth analyses or as a foundation to further subdivide spatial domains and identify novel, data-driven brain region parcellations. However, these spatial domains are based on the 500 genes selected for our panel and do not capture the full diversity of spatially variable genes, leading to an underestimation of regional tissue complexity. This limitation can be addressed by employing genome-wide gene imputation. Our registration workflow achieves high spatial accuracy by precisely aligning cell locations to the Allen Mouse CCFv3 reference framework. This approach produces an improved ABC-atlas dataset with higher-quality cell segmentation and robust anatomical registration, thereby enabling more accurate anatomical and functional analyses. The registration method utilizes clusters of cells with distinct spatial localization as reliable landmarks for effective non-linear alignment. Furthermore, incorporating refined cell type taxonomies into this process can introduce additional spatially specific cell types, which serve as enhanced landmarks and further improve registration accuracy. Furthermore, the registered data facilitate seamless integration with other modalities, including single-cell morphology, patch-seq, and connectivity datasets. This integrative approach enhances the depth and breadth of analyses, allowing for comprehensive characterization of cellular identities and functional properties within anatomically defined regions. Looking ahead, the registration of additional datasets will further increase sampling density, thereby improving the resolution and robustness of spatial transcriptomic studies. Gene imputation allows researchers to investigate the spatial localization of genes not measured in the original experiments. This offers the opportunity to identify region-specific genes or to investigate ligand/receptor interactions. However, caution must be taken when interpreting these results as false positives are easily possible. Imputed gene expression should be used to develop hypotheses but before any conclusions are drawn it is necessary to verify the imputed gene expression independently. Additionally, our pipeline supports scalability and flexibility, making it suitable for large-scale studies and diverse experimental designs. With further optimization, it holds potential for integration into automated systems, streamlining data processing and analysis workflows in various research contexts.

## Methods

### Data

Data generation for mouse and human MERSCOPE data has been described previously (Yao et al., 2023b; Gabitto et al., 2024). Mouse StereoSeq data was obtained from StomicsDB (https://db.cngb.org/stomics/, tissue sections STTS0001453, STTS00014449, and STTS0001457)

### Xenium data generation

Mice were transferred from the vivarium to the procedure room with efforts to minimize stress during transfer. Mice were anesthetized with 0.5% isoflurane. Brains were rapidly dissected and selected based on the absence of dissection damage. The selected brain was flash frozen in OCT using 2-methylbutane chilled with liquid nitrogen and stored at-80°C. The fresh-frozen brain was sectioned at 14 µm using Leica 3050 S cryostats. The OCT block was trimmed in the cryostat until the desired starting section was reached. Sections were collected every 100 µm to evenly sample the brain from posterior to anterior. Fresh frozen tissue sections were mounted onto Xenium slides (10X Genomics) and stored at-80°C. Prior to fixation, slides were equilibrated at 37°C for 1 min using a pre-heated thermal cycler (Bio-Rad 1851197 or Bio-Rad 12015392) equipped with a Xenium Thermocycler Adaptor (10X Genomics 3000954). Fixation was performed by immersion in 10 ml of Fixation Solution (7.5 ml 1× PBS and 2.5 ml paraformaldehyde, Electron Microscopy Sciences 15710) for 30 min at room temperature. Slides were sequentially washed in 1× PBS, 1% SDS (Millipore Sigma 71736), and 70% methanol (Millipore Sigma 34860), followed by additional PBS washes. PBS-T was prepared using 1× PBS and 0.05% Tween-20 (Thermo Fisher Scientific 28320). All solutions were prepared using nuclease-free water (Thermo Fisher Scientific AM9932) and autoclaved glassware. Probe Hybridization Mix was prepared using Xenium Probe Hybridization Buffer (10X Genomics 2000390), Probe Dilution Buffer, and gene expression probes (pre-designed, custom, or standalone). Probes were preheated at 95°C for 2 min and cooled on ice for 1 min prior to mixing. Hybridization was performed overnight (16–24h) at 50°C in a thermal cycler. Slides were washed twice with PBS-T and incubated with Xenium Post Hybridization Wash Buffer (10X Genomics 1000460) at 37°C for 30 min. Ligation was performed using a mix of Xenium Ligation Buffer, Enzyme A, and Enzyme B, prepared fresh and maintained on wet ice. The reaction was incubated at 37°C for 2h. Amplification was carried out using Xenium Amplification Mix and Amplification Enzyme, incubated at 30°C for 2h. Autofluorescence quenching involved sequential incubations with diluted Reducing Agent B, 70% ethanol, and 100% ethanol, followed by incubation with Xenium Autofluorescence Mix (10X Genomics) for 10 min at room temperature in the dark. Slides were rehydrated with PBS and PBS-T, then stained with Xenium Nuclei Staining Buffer for 1 min. After four PBS-T washes, slides were stored at 4°C in the dark until imaging. Slides were loaded into the Xenium Analyzer (10X Genomics 1000481) along with decoding modules, reagent bottles, and consumables. Imaging buffers were freshly prepared, including Xenium Probe Removal Buffer, Sample Wash Buffers A and B, and Instrument Wash Buffer. Imaging was initiated following automated system checks and overview scans of DAPI and autofluorescence. Imaging regions were designated manually, and data acquisition proceeded over 1–3 days. Upon completion, the instrument was cleaned, and slides were post-processed with PBS-T and stored at 4°C. Imaging datasets were previewed for quality control and archived.

### Spots-in-space

To process spatial transcriptomics data, we used Spots-in-Space (SIS) a custom made python package. The package contains three main classes, SpotTable, SegmentedSpotTable, and SegmentationPipeline. SpotTable represents a spatial transcriptomics dataset, containing information about the position of each detected transcript, the associated gene, and image stains. It can be used to manipulate and analyze spatial transcriptomics data. Importantly, it aligns image and transcript data so that one can easily query regions and view both the image and transcript data. SegmentedSpotTable represents a spatial transcriptomics dataset that has been segmented. SegmentationPipeline is the base class for running segmentation on a whole section (or subregion). Its properties are inherited by the MerscopeSegmentationPipeline, XeniumSegmentationPipeline, and StereoSeqSegmentationPipeline classes. For all these classes we provide various utility functions.

### Cell Segmentation

Cell segmentation was performed using Cellpose (Stringer et al., 2021), a generalist deep learning algorithm pretrained on a diverse dataset of 608 images. While the default cyto and nuclei models typically generalize well, they performed poorly on our MERSCOPE data, particularly in densely packed regions. We therefore fine-tuned a Cellpose model using a human-in-the-loop approach (Pachitariu and Stringer, 2022) via the Cellpose GUI, in which model predictions are iteratively corrected by the user and incorporated into training. To generate a robust and representative training set, we selected 9 two-channel 200 × 200 µm images from brains across a range of species and cellular densities: three from human, three from macaque, and three from mouse. Despite aiming for 3D segmentation, training was restricted to the XY plane due to the limited z-resolution and anisotropy in the MERSCOPE images, which rendered XZ and YZ views less informative. Full MERSCOPE images were subdivided into overlapping 350 × 350 µm tiles to alleviate memory limitations and facilitate parallelized processing on a high-performance computing cluster. For cytoplasmic segmentation, we constructed a synthetic intensity image from mRNA transcript locations. Transcripts were binned into a 2D histogram aligned with the DAPI channel and convolved with a Gaussian filter (σ_z = 1, σ_x/y = 3), followed by a 3D median filter (z = 2, x/y = 10), producing a stain-like signal that improved cytoplasmic boundary detection. Segmentation was performed in 3D using Cellpose’s volumetric mode, which computes flows across orthogonal planes (YX, ZX, ZY) and averages them before running 3D dynamics. Although MERSCOPE data contained only 7 z-planes, this method outperformed 2D segmentation followed by stitching. Rather than merging segmentation masks across the entire section, we assigned transcripts to segmented cells within each tile and subsequently merged cell IDs. Transcripts appearing in overlapping regions were reconciled: if >50% of a smaller cell’s transcripts were shared with a larger one, the cells were merged; otherwise, the transcript was assigned to the newer cell during tile stitching. To represent cell boundaries, we constructed 2D alpha shapes of transcript positions on each z-plane. The alpha parameter was defined as the maximum value producing a single closed polygon and scaled by 0.75 to enforce biologically realistic shapes. These planar polygons were stacked to generate a pseudo-3D representation of each cell. All resulting data were stored in an AnnData object with cell-by-gene matrices and associated metadata. The structure of the output is summarized below:

### Data

- X Cell-by-gene matrix of segmented transcript counts obs: Cell-level metadata:
- Transcript centroid
- Polygon centroid (area weighted)
- Volume (3D)
- Area (2D)
- Original cell ID
- Production cell ID (globally unique)

var: Gene-level metadata

- Gene name
- Number of cells expressing each gene
- Total number of segmented transcripts
- Total unsegmented transcripts

uns: Unstructured metadata

- Git version of sis repository
- Cell polygons (GeoJSON FeatureCollection)
- Segmentation parameters

### Identification of incongruent genes

To identify cluster-specific marker genes from the single-cell RNA sequencing (scRNA-seq) data, we performed a differential expression analysis using the Scanpy Python package (Wolf et al., 2018). The analysis was run on the previously generated log-normalized expression matrix. To decrease computational demand, we did not run the analysis on the original single-cell dataset with about 7 million cells but instead used a gene expression matrix of the cluster medians. We utilized the sc.tl.rank_genes_groups function to find genes enriched in each cell cluster compared to all other cells (one-vs-rest strategy). The Wilcoxon Rank-Sum test (method=‘wilcoxon’) was selected for this purpose. We picked the top genes for each division.

### Doublet removal

To identify doublets, we applied the Solo algorithm (Bernstein et al., 2020) independently to each section. Solo is a semi-supervised deep learning method that embeds cells using a variational autoencoder and subsequently trains a feed-forward neural network to classify observed cells against simulated doublets. Solo outputs a probability score for each cell being either a singlet or doublet. We computed the difference (dif) between the singlet and doublet scores of the predicted doublets. To define a threshold for doublet classification, we calculated the 0.9 and 0.1 quantiles (q) of the dif distribution among the predicted doublets. The threshold was set to q0.9(dif) - q0.1(dif), and all cells with dif values above this threshold were classified as doublets. The threshold was determined by sampling sections and subregions along the anteroposterior axis and evaluating false-positive and false-negative doublets in each case. These cells were excluded from the dataset.

### Cell filtering following label transfer

To assign transcriptomic cell identities, we used MapMyCells (MMC), a bootstrapped, correlation-based mapping algorithm as described in Yao et al. (2023b). Each cell is assigned a correlation score for every reference cell type, and the label with the highest correlation is used as the predicted identity. Previously, cells with a top correlation score below 0.5 were excluded. However, this global threshold does not account for variability in the correlation score distributions across different cell types. To adaptively threshold cell quality based on the distribution of mapping scores for each target cell type, we employed the Median Absolute Deviation (MAD) and extended it to a DoubleMAD approach to better accommodate asymmetric or heavy-tailed distributions. MAD is defined as the median of the absolute deviations from the median of a distribution. In standard implementations, this single measure is applied symmetrically to both tails. However, for skewed distributions—frequently observed in MMC mapping results—this can fail to identify low-quality cells in the lower tail. In the DoubleMAD framework, we compute separate deviations above and below the median: MAD^low^: median absolute deviation of values below the median MAD^high^: median absolute deviation of values above the median. Cells are flagged as low-quality if their top correlation score falls below the threshold: Median – 3 × MAD^low^ For bimodal distributions, we refined this procedure to avoid penalizing cells in the lower mode when both modes represented biologically plausible identities. Bimodality was assessed using the LaplacesDemon R package, and a distribution was considered bimodal if it meets all of the following criteria: The smaller mode contained at least 10% of the total density. A local minimum was located between median – 0.05 and median + 0.05. The difference between the two mode peaks was less than 0.05 When a cell type’s correlation distribution was classified as bimodal, we isolated the portion of the distribution above the local minimum and applied the DoubleMAD filter to that subset only. This ensured that the filter did not erroneously discard a biologically valid subpopulation. Only cells that passed initial quality control thresholds for gene count, transcript count, proportion of unassigned (“blank”) transcripts, and doublet detection were included in the MMC filtering step.

### Spatial domain detection

To identify spatial transcriptomic domains, we implemented a multi-step computational pipeline. First, raw spot-level gene expression data were downsampled by aggregating them into 30 µm spatial grids. Quality control (QC) filtering was applied to retain grids containing at least 60 detected genes, 300 transcripts, and less than 3% blank values. Following QC, we applied STAligner (Zhou and Zhang, 2023), a spatial embedding model that constructs a spatial neighborhood graph using a k-nearest neighbors (KNN) approach (k = 8), to encode local spatial relationships between spots. Spatial embeddings generated by STAligner were clustered using the Leiden algorithm across a range of resolution parameters (0.6–2.0) to identify transcriptionally coherent domains. To ensure anatomical consistency across serial sections, Scube was used to infer and apply rigid transformations that rotate individual tissue sections, aligning them along a common superior-inferior axis. This step effectively reconstructs a pseudo-3D spatial volume from 2D sections and facilitates downstream comparative analyses across the tissue stack. STAligner is a graph-based deep learning framework designed to learn spatially informed embeddings that integrate local gene expression patterns with spatial context across sections. The method consists of four key steps:

1. **Spatial Graph Construction:** Gene expression matrices were normalized, and a spatial neighbor graph was constructed using spot coordinates to model local spatial structure. The graph topology can be defined using either a k-nearest neighbors (KNN) or radius-based method; in this study, we used a KNN graph with 8 neighbors per node.
2. **Spatial Embedding Learning:** Embeddings were learned via a graph-based autoencoder trained to encode both transcriptomic and spatial features, producing low-dimensional latent representations that preserve neighborhood relationships.
3. **Triplet Loss for Cross-Section Alignment:** To align embeddings across serial tissue sections, a triplet loss objective was employed. Each triplet comprised of an anchor spot, a positive (a mutual nearest neighbor from a different section with similar gene expression), and a negative (a spatially distant spot from the same section with dissimilar expression). The loss was designed to minimize the distance between anchor-positive pairs and maximize the distance to anchor-negative pairs, thus encouraging alignment of biologically similar regions across sections.
4. **Iterative Triplet Selection and Training:** The triplet construction and embedding optimization were performed iteratively. As embeddings evolved during training, triplet relationships were updated to reflect the current structure of the embedding space, allowing for progressive refinement of spatial alignment.

Clustering was performed using the Leiden algorithm implemented in the RAPIDS single-cell library. A KNN graph was constructed using 15 neighbors based on the STAligner embedding stored in the AnnData object. Bootstrap-based cluster stability analysis was performed by subsampling 80% of cells from the dataset 25 times and performing Leiden clustering at resolutions for 0.2 to 2.0 using the STAligner embedding. Cluster stability was quantified by calculating pairwise Adjusted Mutual Information (AMI) and Adjusted Rand Index (ARI) scores. The number of clusters identified in each iteration was recorded to assess clustering consistency. We settled on a final resolution of 1.4. STAligner embeddings were visualized as UMAP and colored by cluster assignment.

### Image Registration

To minimize manual alignment, we used sequential tissue sections that were internally rotated relative to each other as generated by Scube (Xu et al., 2023). Images of cell centroid locations were produced and colored according to their Common Coordinate Framework (CCF) region assignments. These images were ordered by relative anterior–posterior (A–P) position and imported into QuickNII via the FileBuilder tool, using the Rainbow 2017 reference atlas for initial rigid alignment. Sections with clear anatomical landmarks were manually adjusted to determine their precise A–P position within the CCF, while the positions of intervening sections were interpolated. For the selected reference sections, rotation, tilt, and scaling were refined to match the CCF template, and alignment was iteratively optimized across the full series. Final QuickNII registrations were saved in both XML format (for potential future adjustment in QuickNII) and JSON format (for subsequent processing in VisuAlign). QuickNII transformations were imported into VisuAlign, where anatomical anchors were placed on corresponding landmarks and adjusted to match visible structures in the tissue sections. This procedure was applied consistently across all landmarks in the dataset. Two-dimensional tissue coordinates were then transformed into 3D CCF coordinates. Landmark points defined in the VisuAlign JSON – containing both source and target user-corrected marker positions – were first used to perform a Delaunay triangulation of each section. Cell coordinates were scaled to match the input image resolution, then mapped to the triangulated mesh created from the landmark points. For each cell, barycentric coordinates – weights describing its relative position with respect to the triangle’s three vertices – were calculated to interpolate the cell’s corrected position. Coordinates were then normalized to image dimensions and transformed using the 3×3 affine transformation matrix stored in the QuickNII JSON. Finally, the resulting QuickNII 3D coordinates were multiplied by the Allen Reference Atlas (ARA) transformation matrix to scale them to voxel size and align them to the ARA coordinate system origin. Brain region annotations were assigned by mapping each cell’s CCF coordinates to the 25 µm ARA annotation volume.

### Gene Imputation

To address the gene panel limitations of the MER-SCOPE experiment, we performed gene imputation using ENVI (Haviv et al., 2025), a conditional variational autoencoder that integrates scRNAseq and spatial transcriptomics data into a shared latent space. As a prerequisite, we applied the covariance environment (COVET) framework to represent spatial neighborhoods. In COVET, the niche of each cell is defined as its k nearest spatial neighbors (k = 8) and summarized as a gene-gene shifted covariance matrix. Unlike mean expression, covariance captures relationships among genes and cell states shaped by local interactions, providing a robust and information-rich representation of the niche. The shifted formulation normalizes covariance relative to the global dataset mean, enabling direct comparison across niches and increasing robustness to technical artifacts. This per-cell COVET representation was then used as input to ENVI to provide spatial context. Spatial cell counts were encoded into a 512-dimensional latent vector using the ENVI encoder, and the resulting latent representations were passed through the expression decoder with the scRNAseq auxiliary variable to predict the full transcriptome for each spatial cell. The model was trained jointly on both single-cell and spatial datasets, using a negative binomial likelihood for single-cell counts and a Poisson likelihood for spatial counts, with KL divergence regularization to constrain the latent space. This approach leverages transcriptome-wide information from scRNAseq data to recover unmeasured genes in the spatial dataset while accounting for modality-specific biases. For ENVI application, the MERSCOPE dataset was first divided into major brain regions (Isocortex, OLF, HIP, RHP, CTXsp, STRd, STRv, LSX, sAMY, PAL, HY, TH, MB, P, MY, CB, VS, and fiber tracts), and datasets were prepared for individual sections or hemisections. Because ENVI requires both spatial and scRNAseq modalities, an equal number of cells were selected from the Allen Brain Cell (ABC) atlas by matching the corresponding regions. Subsampling of scRNAseq cells was performed using the scValue Python package (Huang et al., 2025), which trains a random forest classifier to assign a “data value” to each cell based on its importance in distinguishing cell types. Cells with the highest values were selected using a full-binning strategy, resulting in a reduced dataset enriched for the most informative cells. For VS and fiber tract regions, which lacked direct ABC atlas counterparts, cells were selected from the ABC atlas based on the cell types present in the VS or fiber tract regions of the processed section, taking only cells that belonged to ABC atlas regions included in the processed section (i.e., regions outside VS and fiber tracts). After subsampling, both spatial and scRNAseq datasets were provided as input to ENVI to learn a shared latent space and perform gene imputation.

### Assessing imputation accuracy

To test the imputation accuracy, we used an unpublished dataset that was imaged with a gene panel different from the one employed in this study. In total, 263 genes from this panel were not present in the studied dataset. We aligned hemisections from the imputed dataset and the datasets with the alternative panel (24 hemisections) located at approximately the same antero-posterior level using the CAST algorithm (Tang et al., 2024). Briefly, CAST builds a tissue graph from spatial coordinates and gene expression data, learns its structure with a graph neural network, and applies rigid and non-rigid transformations to align tissue sections across datasets while preserving spatial relationships. After alignment, we assessed imputation accuracy with the Multiscale Spectral Similarity Index (MSSI) method described in Haviv et al. (2025). In brief, MSSI compares expression profiles of two genes from a multiplexed image, either two distinct genes or a ground-truth gene and its imputed value, while also incorporating spatial coordinates. A k-nearest neighbor (k = 8) graph is first constructed from spatial coordinates, followed by iterative graph coarsening. This process pools nodes based on connectivity patterns in a manner analogous to image downsampling. The graph is coarsened and blurred four times by a factor of two, producing multiscale representations of gene expression that are compared using a structural similarity index based on luminance, contrast, and correlation components. In this study, we modified the MSSI framework to account for differences in dataset size and spatial resolution. Specifically, (i) we allowed for datasets with non-matching spatial coordinates and different numbers of cells, (ii) we applied separate coarsening operators for reference and query datasets, ensuring that each graph was downsampled according to its own geometry, and (iii) we replaced direct cell-to-cell comparisons with a distribution-based similarity function that normalizes expression values, computes luminance and contrast similarity, and evaluates structural similarity via correlations between histograms of the expression distributions. These modifications enable MSSI to robustly compare aligned datasets that do not share the same cellular segmentation while maintaining the multiscale framework of the original method. Thus, we computed the MSSI for the 263 imputed genes absent from the studied dataset and compared them to the expression profiles measured in the two datasets with original MERFISH expression.

## Acknowledgments

We are grateful to the Transgenic Colony Management, Lab Animal Services, Spatial Transcriptomics, and Histology teams at the Allen Institute for technical support. The research was funded by the U19MH114830 grant from National Institute of Mental Health to H.Z, under the BRAIN Initiative of the National Institutes of Health (NIH). The content is solely the responsibility of the authors and does not necessarily represent the official views of the NIH and its subsidiary institutes. This work was also supported by the Allen Institute for Brain Science. The authors thank the Allen Institute founder, Paul G. Allen, for his vision, encouragement, and support.

## Data and code availability

Segmentation pipeline can be found at https://github.com/AllenInstitute/spots-in-space. The processing pipeline can be found at https://github.com/AllenInstitute/Spatial-Transcriptomics-Processing-Pipeline.

