## Supplementary Information for "SCALPEL: A pipeline for processing large-scale spatial transcriptomics data"

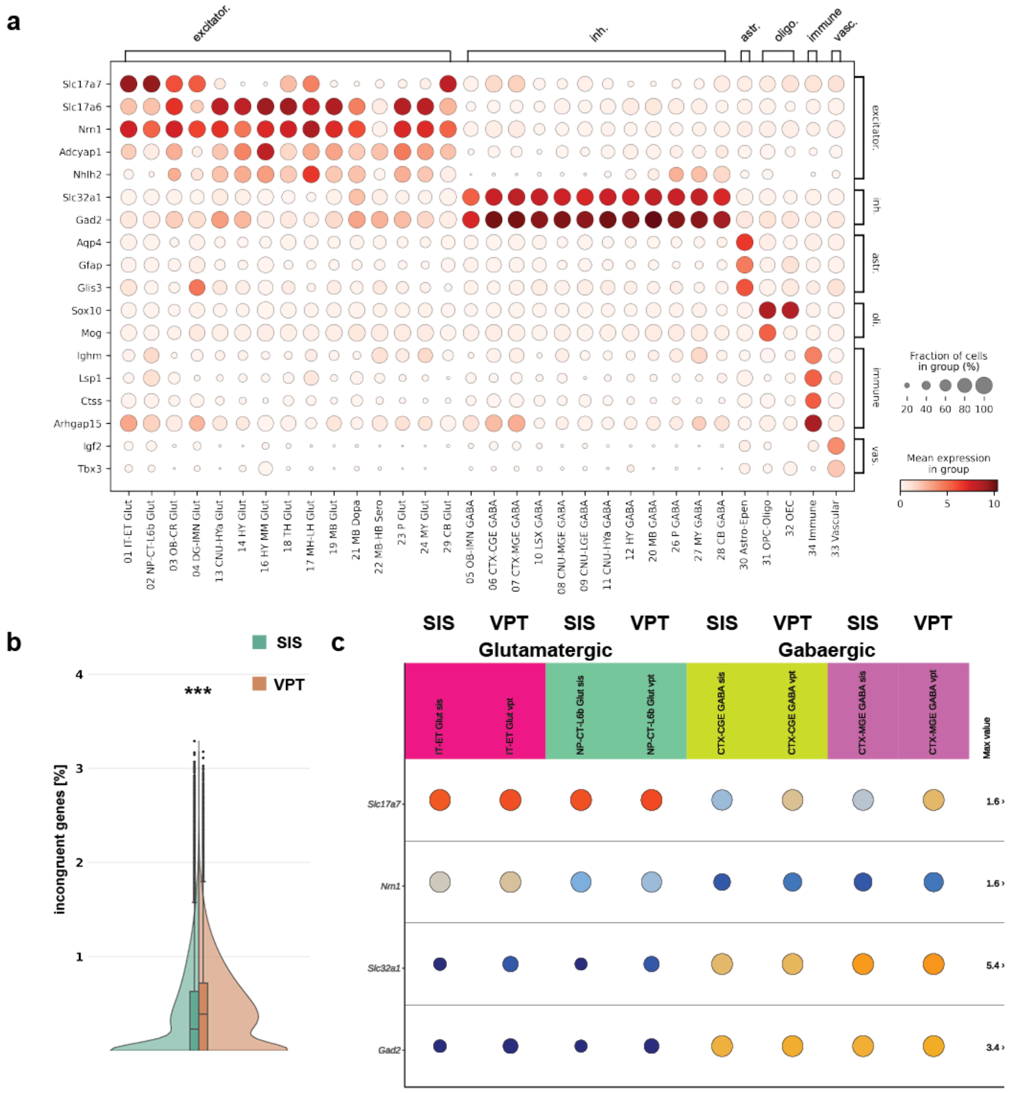


**Figure S1. Incongruent genes.**
(**a**) Dotplot showing the expression of incongruent genes at the division level. (**b**) Distribution of percentage of incongruent genes in SIS (green hue) and VPT (orange hue) segmented cells. c). Dotplot comparing expression of marker genes for glutamatergic (Slc17a1 and Nrn1) and GABAergic (Slc32a1 and Gad2) cells between SIS and VPT cells. (**d**-**f**) Comparison of cell volume distributions for selected subclasses with known size disparities.

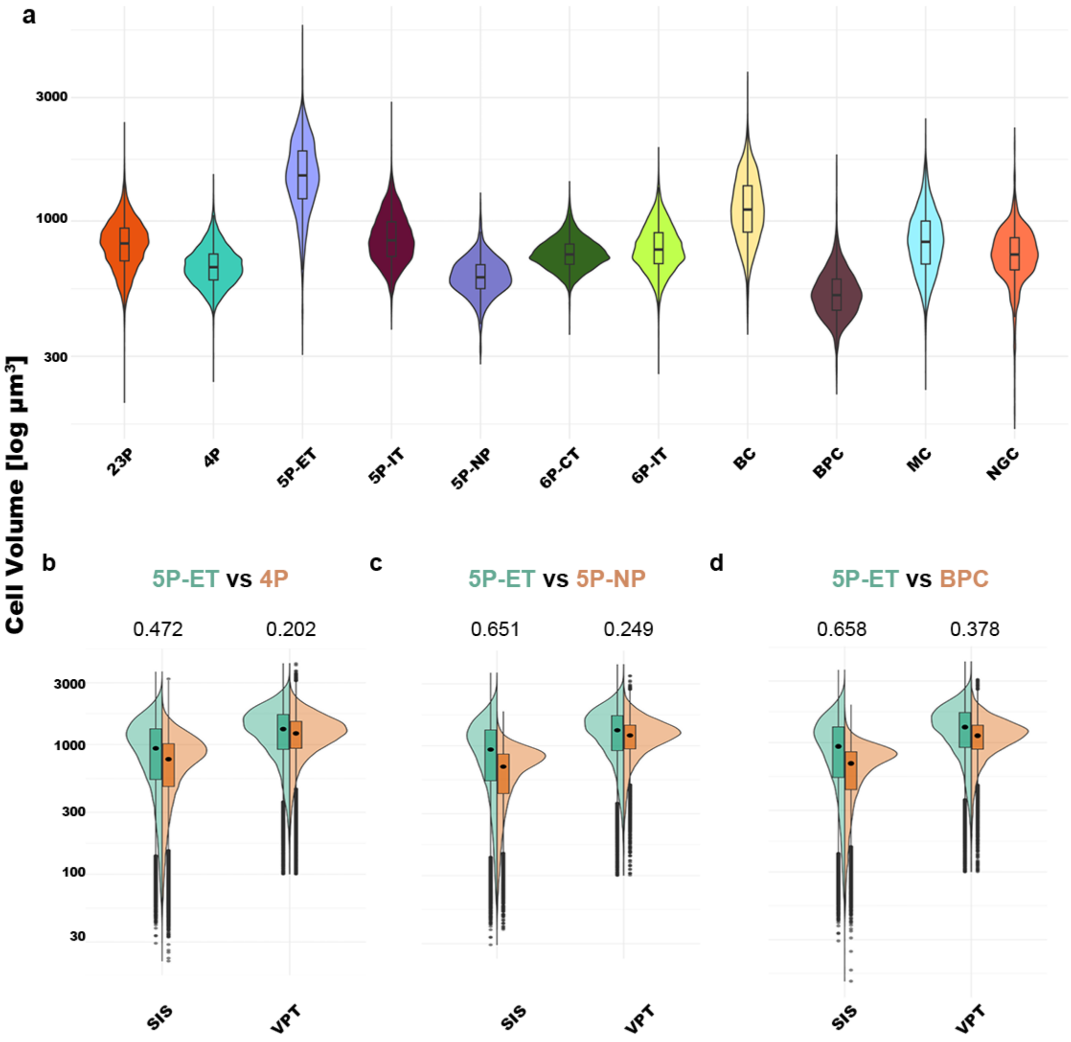


**Figure S2. Cell volume comparison.**
(**a**) Violin plots showing distribution of cell volumes for predicted subclasses in reference EM dataset. (**b**-**d**) Comparison of cell volumes of select subclasses from MERSCOPE data segmented using either (green hue) or VPT (orange hue). Values above each split violin plot indicate Cohen's D.

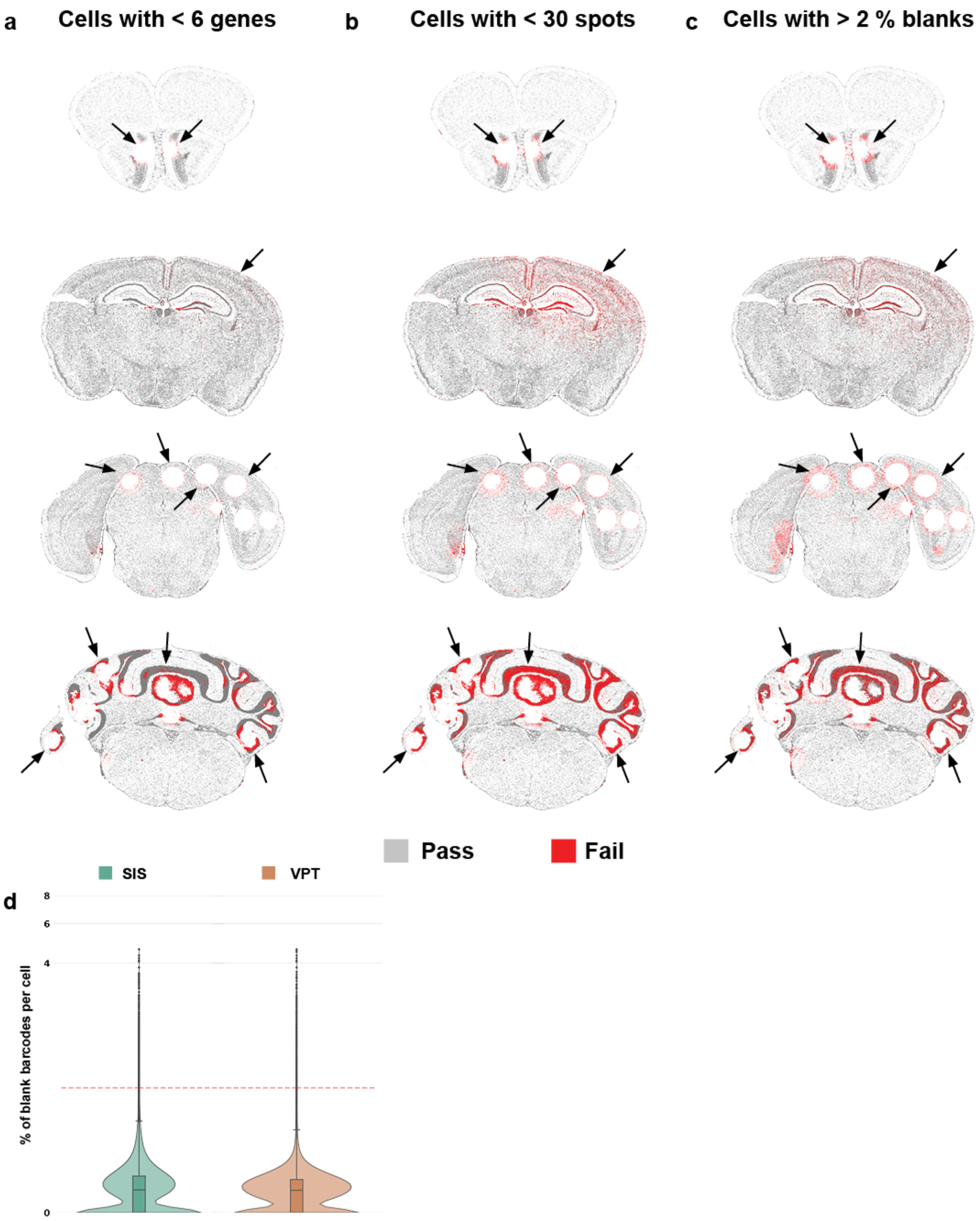


**Figure S3. Low-quality cell filtering threshold.**
(**a**-**c**) Example section highlighting the spatial location of cell filtered based on low gene detection (a), low transcript detection (b), and high percentage of blank barcodes (c). (**d**) Violin plots showing the distribution for percentage of blank barcodes between SIS (green hue) and VPT (orange hue) segmented cells.

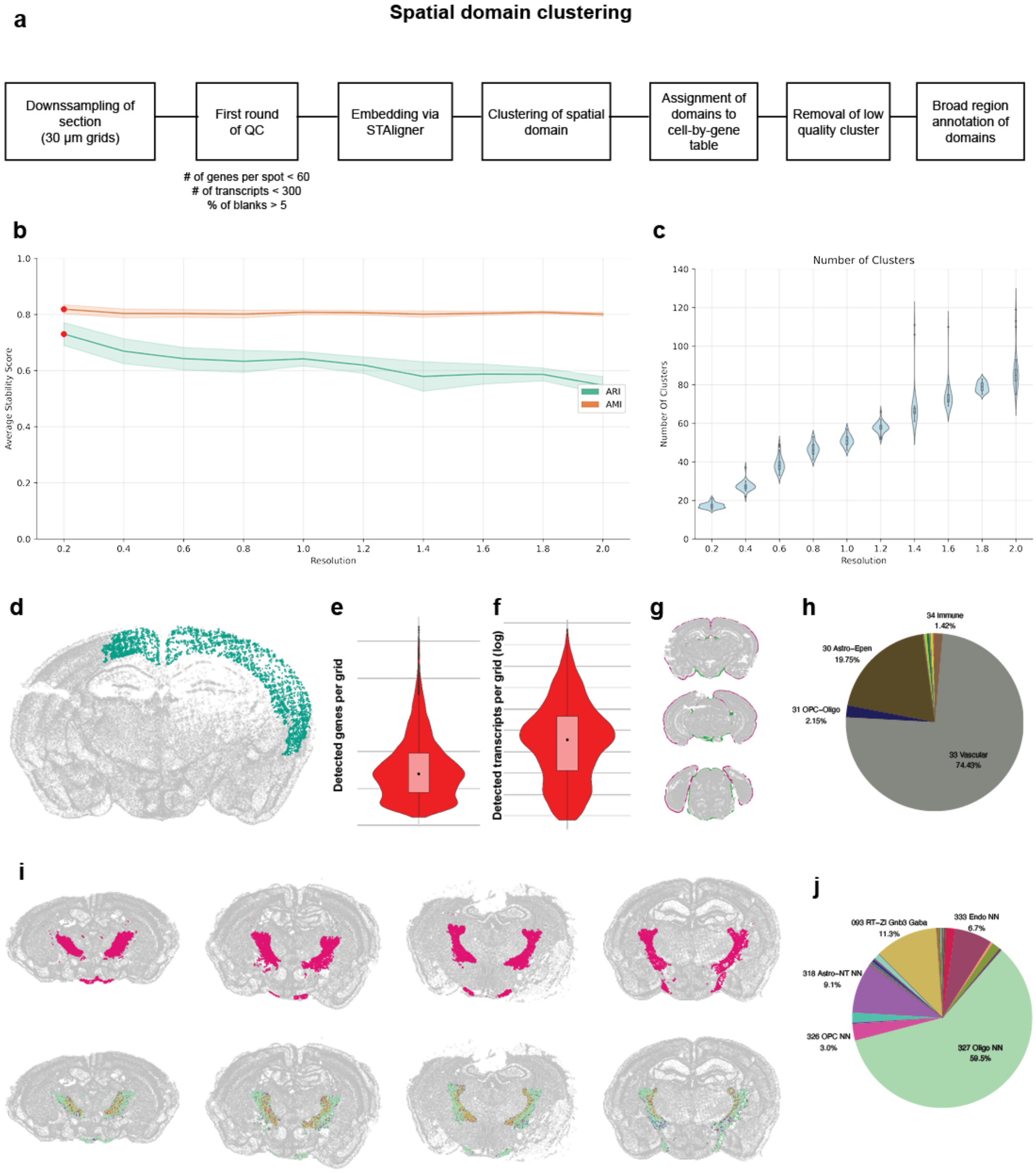


**Figure S4. Evaluation of spatial domains.**
(**a**) Schematic of workflow for processing spatial domain data. (**b**) Adjusted Rand Index (ARI) and Normalized Mutual Information (NMI) between Leiden clusters for various resolutions. (**c**) Distribution of number of clusters derived from bootstrapped Leiden clustering at various resolutions. (**d**) Example section for low-quality cluster (green). (**e**-**f**) Violin plot showing distribution of gene (e) or transcripts (f) per grid for low quality cluster from d. (**g**) Example sections showing spatial domains Borders_1 (purple) and Borders_2 (green). (**h**) Pie chart showing the distribution of subclasses within spatial domains Borders_1 and Borders_2. (**i**) Upper panel: Representative section showing the spatial domain fiber_tracts_3. Lower panel: Same sections as in upper panel but spatial domain is colored by subclass. (**j**) Pie chart showing the distribution of subclasses within spatial domain fiber_tracts_3.

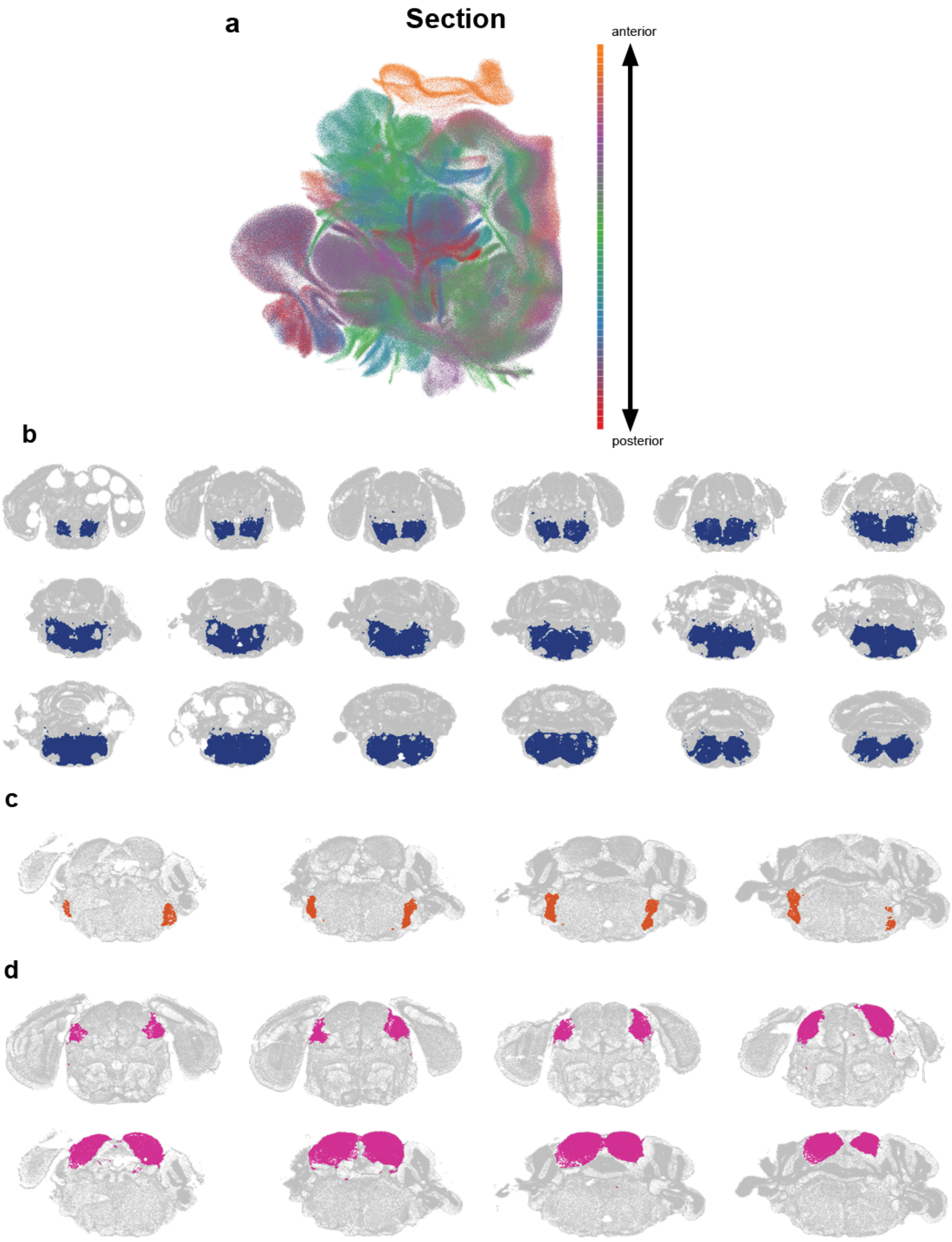


**Figure S5. Evaluation of spatial domains.**
(**a**) UMAP of STAligner embedding colored by relative position of section along the anterior to posterior extent of the brain. (**b**-**d**) Example sections showing the localization of spatial domains P-MY_1 (b), PSV_1 (c), and IC_1 (d).

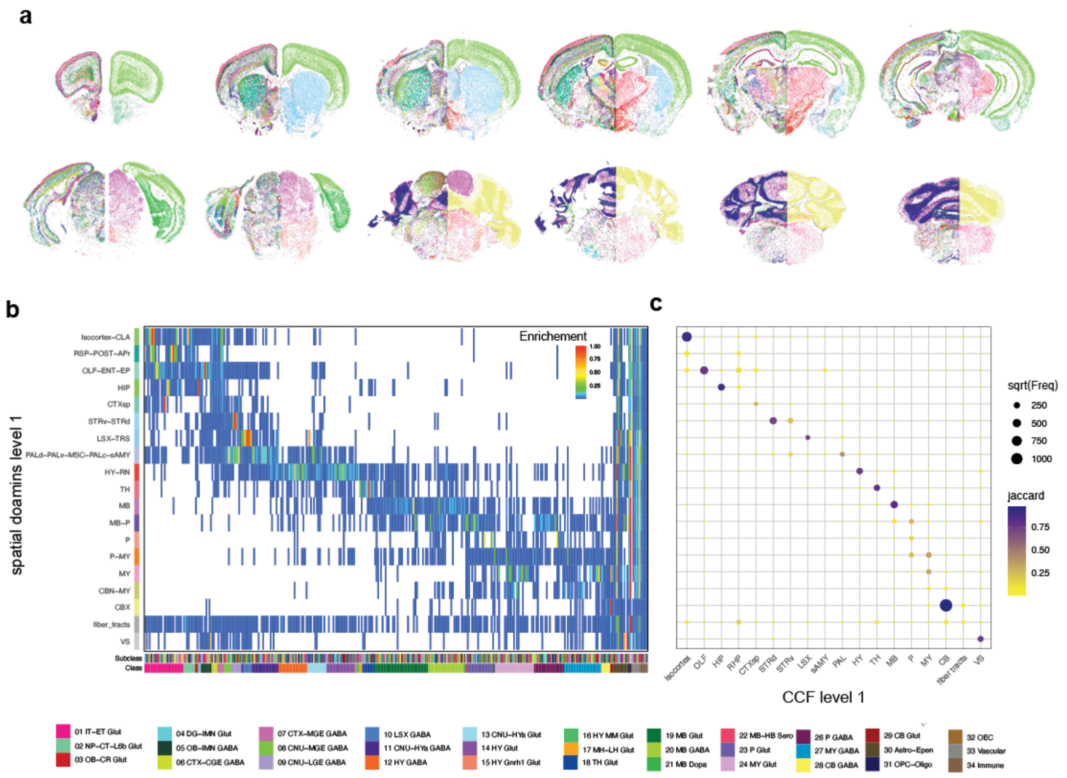


**Figure S6. Broad region specific clusters.**
(**a**) Example section highlighting broad region assignment of clusters. Left half of each section is labeled by cell cluster and right half based on broad region to which it was manually assigned. (**b**) Heatmap showing subclass enrichment in spatial domains aggregated by CCF regions. (**c**) Dotplot demonstrating the Jaccard overlap of region-specific cell types in spatial domains aggregated by CCF.

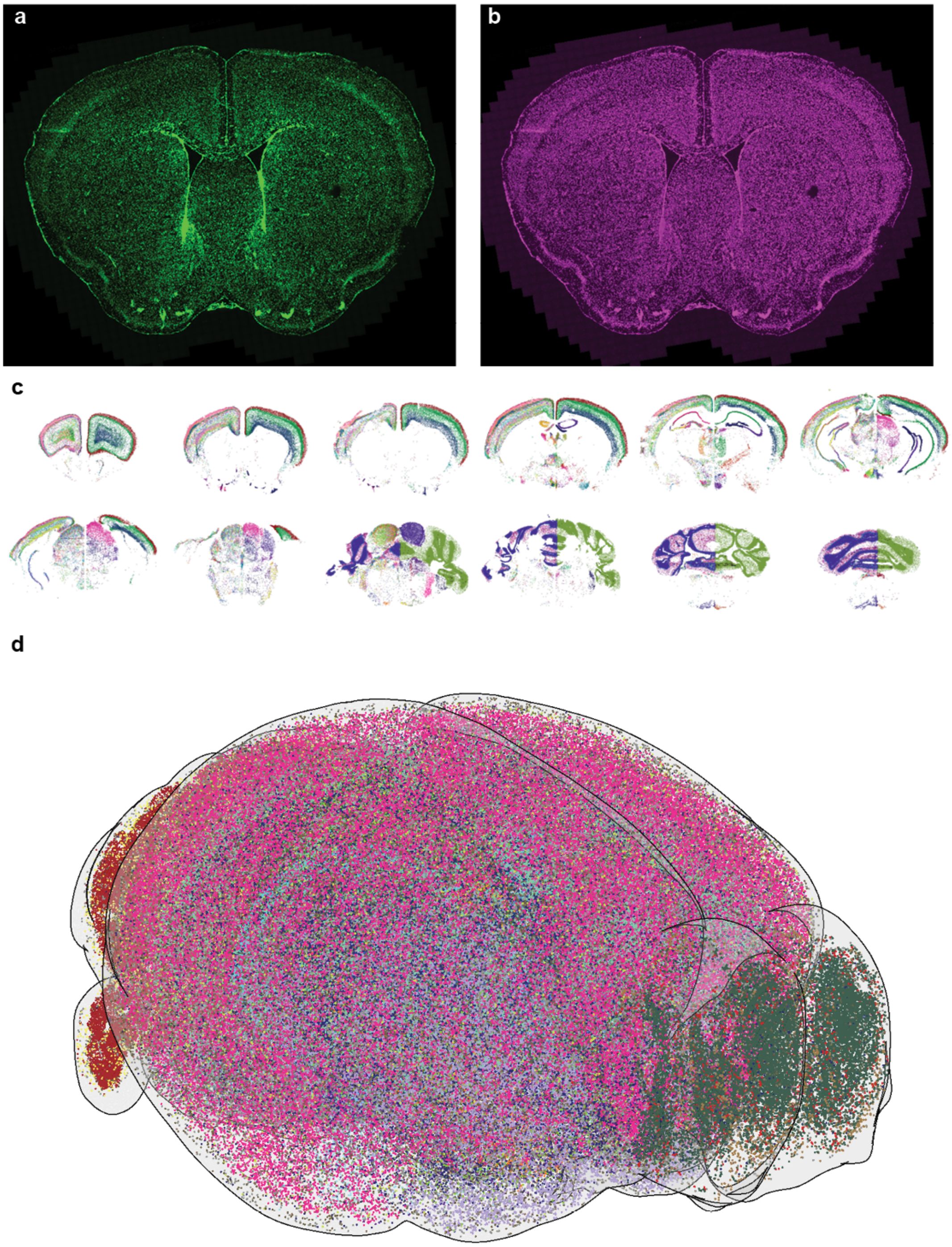


**Figure S7. Registration.**
(**a** and **b**) Images of example sections with staining for DAPI (a) and PolyT (b). (**c**) Example section highlighting landmark assignment of clusters. Left half of each section is labeled by cell cluster and right half based on landmark to which it was manually assigned. (**d**) 3D view of cells labeled by classes in CCFv3. For computational purpose, the total cells were subsampled to 30%.

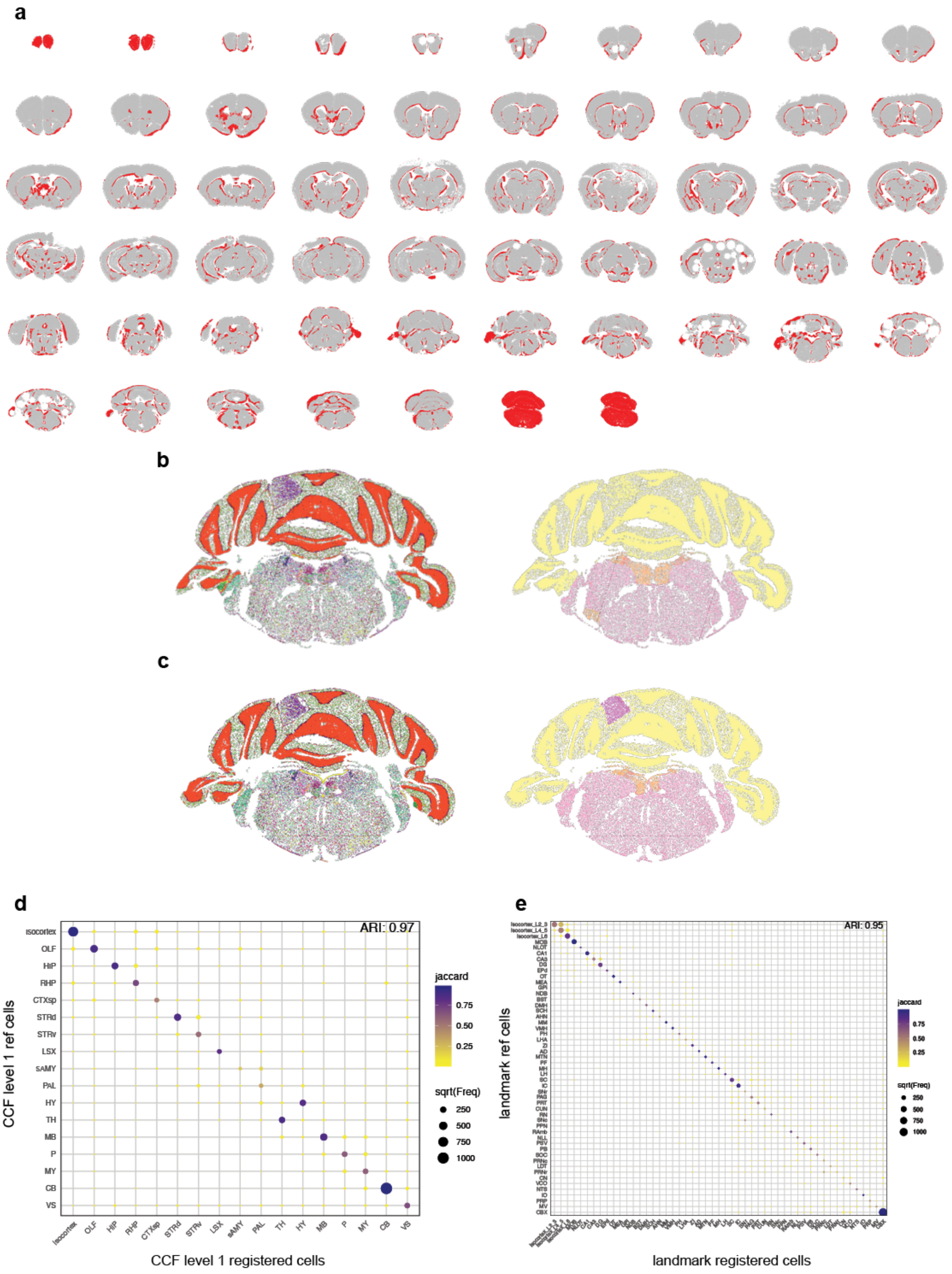


**Figure S8. Registration evaluation.**

(**a**) Overview of all sections indicating newly added cells in red. (**b**) Old registration results. Left-hand side shows example section labeled by subclass, right-hand side shows same section labeled by broad region CCF assignment. (**c**) New registration results. Left-hand side shows example section labeled by subclass, right-hand side shows same section labeled by broad region CCF assignment. (**d**) Dotplot highlighting the overlap of broad specific cells and their CCF region assignment using old registration. Size indicates number of cells and color Jaccard coefficient. ARI = adjusted rand index. (**e**) Same as in d but instead of broad regions more fine grain landmarks are compared.

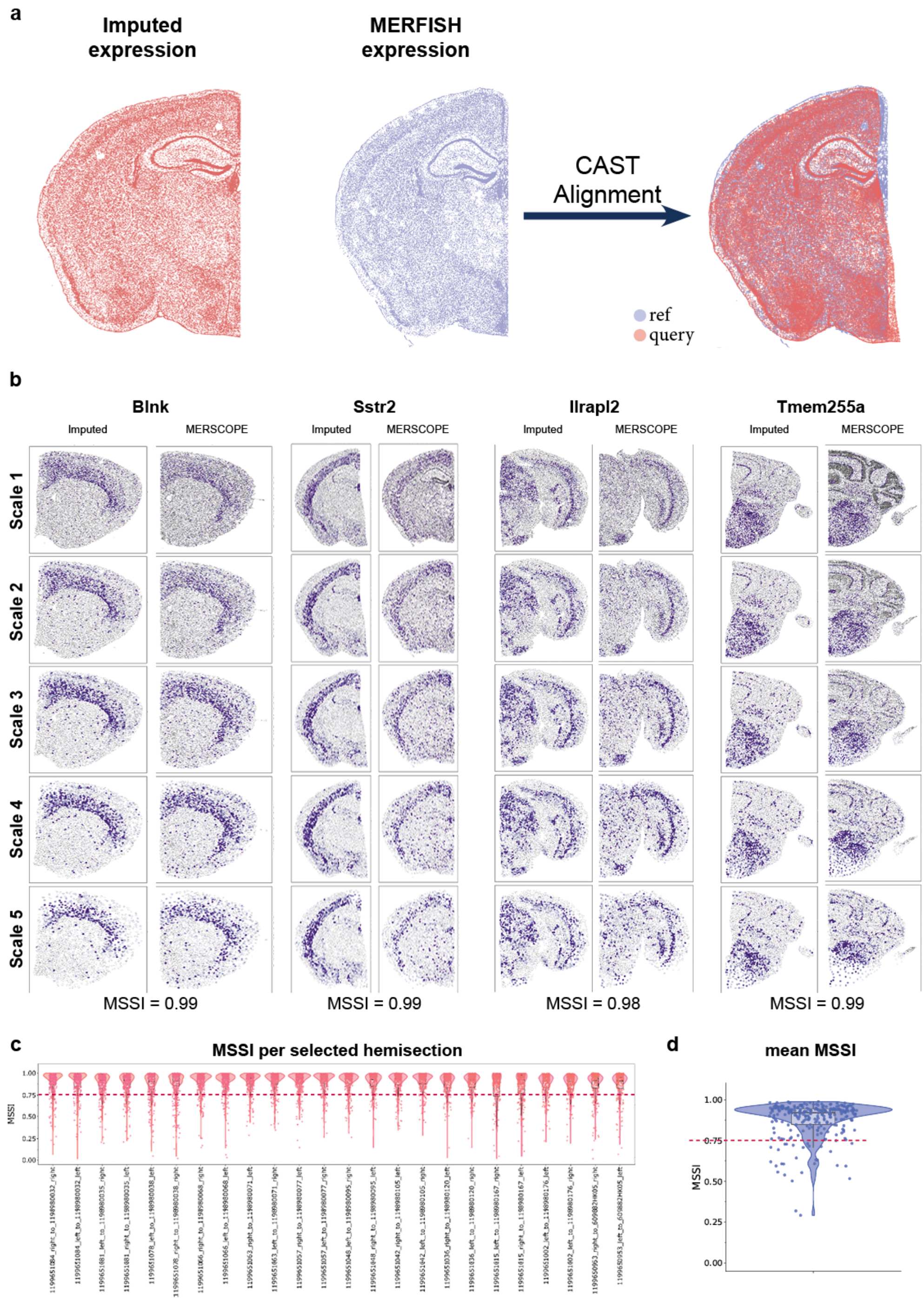


**Figure S9. Imputation evaluation.**
(**a**) To compare sections from two different experiments using the Multiscale Spectral Similarity Index (MSSI), we identified the corresponding sections from the two experiments and aligned the query section (imputed gene expression) to the reference section (MERFISH expression) using CAST. (**b**) Representative examples of imputed genes with high MSSI. Left column is imputed gene expression and right column is original MERFISH expression. Each row is the spatial gene expression at a different resolution. (**c**) Violin plot showing the distribution of MSSI for all 263 genes in each section pair. (**d**) Violin plot showing the distribution of the mean MSSI for all 263 genes.

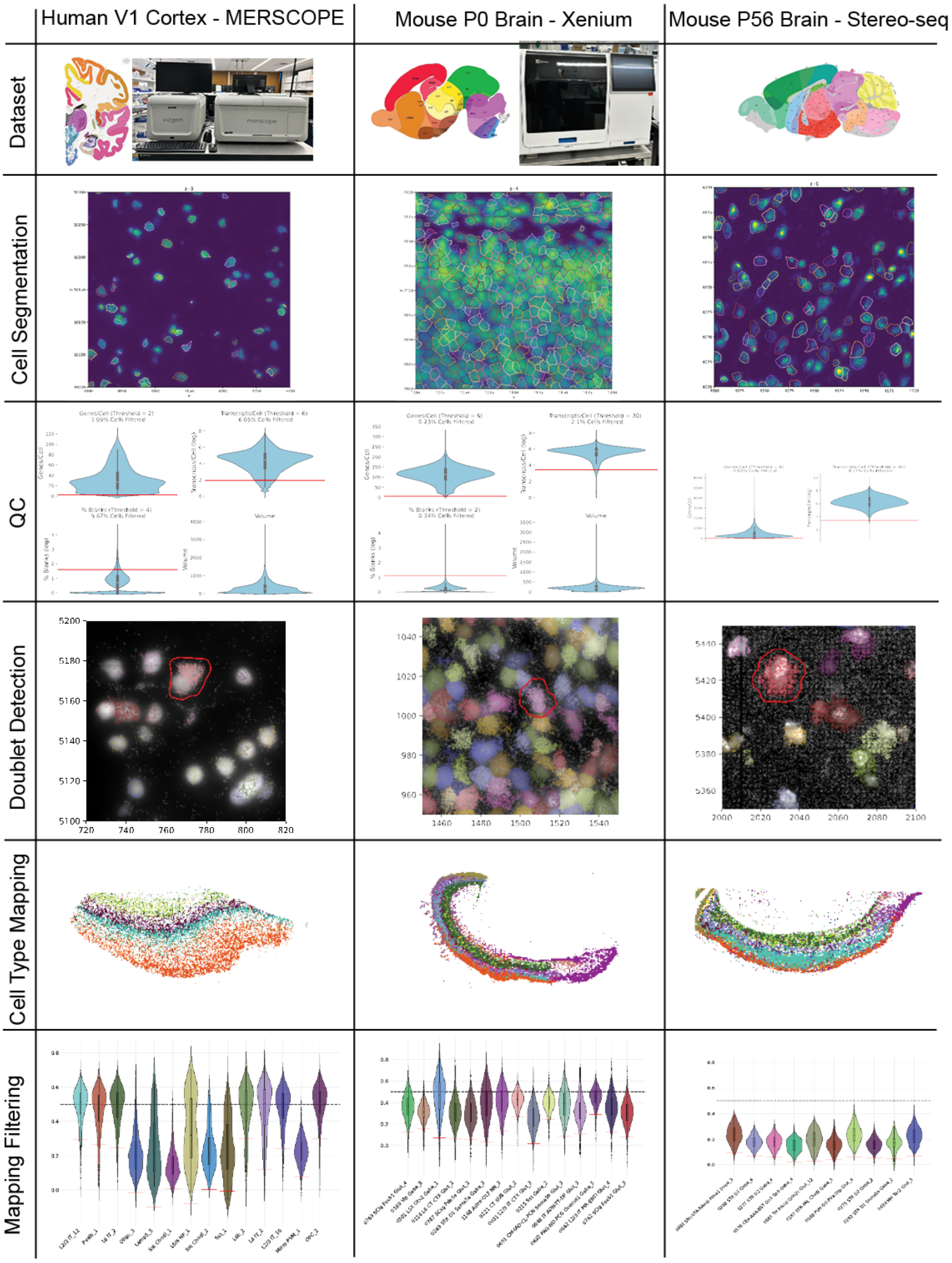


**Figure S10. Processing pipeline for different datasets. .**
We used our basic processing pipeline on human MERSCOPE data (left), P0 mouse brain Xenium data (middle) and adult mouse StereoSeq data (right).
